# How the rhizosphere chemistry explains the effectiveness of radish in reclaiming legacy phosphorus compared to maize

**DOI:** 10.1101/2025.04.06.647426

**Authors:** Yahaya Mohammad Yusuf, Gabriel Sgarbiero Montanha, Kaoutar Benghzial, Paulo S. Pavinato, Hudson Wallace Pereira de Carvalho

## Abstract

**Background and aims:** Around 50% of phosphorus (P) applied to tropical soils is not used by plants and become part of the legacy P. Some cover crops can extract this P. But how do they do this and which form do they extract most?

**Methods:** Radish (*Raphanus sativus* L.) and maize (*Zea mays* L.) were planted in rhizoboxes and tubes containing a mixture of sand, kaolinite, hematite, and boehmite, the later and former bearing sorbed phosphate. The structure and composition of the root-soil interface were determined by scanning electron microscopy (SEM) and Fourier-transform infrared spectroscopy (FTIR). Additionally, synchrotron-based microprobe X-ray fluorescence (µXRF) was employed to assess the spatial distribution of Al, Fe, and P. The P speciation at the rhizosphere was evaluated using microprobe X-ray absorption near edge structure (µXANES).

**Results:** The chemical images revealed that both plants depleted more of the Al-bound P than Fe-bound P, with radish demonstrated a higher efficiency compared to maize. The total P uptake by radish from Al-bound P was 42% higher than that uptake from Fe-bound P. Additionally, radish absorbed 34% to 90% more total P compared to maize, indicating a significant difference between the two crops. The superior capacity exhibited by radish seems to be connected to organic acids and total carbon exudation, which the latter was 2.14-fold more than maize.

**Conclusion:** Radish uptakes greater Al-bound P than Fe-bound P. This insight could help in proper management of soils where P is predominantly bound to Al and significantly increases P use efficiency.

## Introduction

Phosphorus (P) is an essential element that plants require in substantial amounts for their development (Hu et al. 2020) and plays a crucial role in the production of food, feed, fiber, and biofuel (Pavinato et al. 2020). Due to the indispensable role of P in agriculture, farmers consistently apply significant amounts of fertilizers to meet the crops’ P requirement (Roswall et al. 2021). Phosphorus use efficiency (PUE) widely reported in the literature is generally low, leading to discussions about the sustainability and economic losses associated with P fertilization. However, the percentage reported varies slightly across countries, this might be influenced by myriads of soils, climate, agronomic and environmental factors. For instance, a recent study revealed that the average PUE in China was 40% between 2005 and 2018 (Shen et al. 2023).

In contrast, Pavinato et al. (2020) reported that PUE in Brazilian croplands averaged approximately 50% over the period from 1967 to 2016. Globally, the average PUE by crop is less than 30% (Benício 2022), because over time, most of the P applied especially to highly weathered soils undergo processes like fixation by soil minerals such as iron (Fe), aluminum (Al) oxyhydroxide and calcium carbonate, precipitation, and complexation with organic matter (Doydora et al. 2020), then transformed into soil P species which are little or even unavailable for plants (Soltangheisi et al. 2020). These phenomena have led to the accumulation of P (legacy P) in some soils to concentrations that are occasionally much higher than the agronomic optimum (Gamble et al. 2020; Wieczorek et al. 2022). Under acidic conditions, most of P is bound to Fe or Al, whereas calcium-bound P is mostly found under alkaline conditions (Shariatmadari et al. 2007).

The buildup of P content in the soil poses a significant challenge, as it may lead to the significant movement of P from soil to water bodies, causing toxic algal blooms and threatening agricultural efficiency (Roswall et al. 2021). For instance, according to Roy et al. (2016), accumulated P residual from past fertilization, has revealed that billions of dollars are deposited in the soil annually. It is estimated that global cropland with high P-fixing potential soils covers about 118−188 million ha, and in a future scenario, this number is expected to increase to about 169−365 million ha, while about 2–8 million tons of P fertilizer are applied annually on this cropland, with half of this fertilizer becoming fixed in the soil (Roy et al. 2016). Thus, more efficient phosphate management is required globally to effectively slow down the depletion of non-renewable natural resources and finite reserves of easily recoverable rock phosphate, as well as ensure its sustainable utilization (Cordell et al. 2009). However, the bioavailability of P fixed by soil minerals varies significantly as a function of both P chemical forms and plant species (Laan et al. 2023; Zhao et al. 2023).

Therefore, implementing practices that lead to the plant utilization of the legacy P is required towards higher PUE (Doydora et al. 2020). Hence, selecting plant species or cultivars with improved ability to reclaim the legacy P would be an important option for managing soils with low P bioavailability (Zhu et al. 2002). Several studies have indicated that the incorporation of cover crops is a promising approach, as they possess strategies for mobilizing legacy P. Hallama et al. (2019) study shows that several cover crops such as those from the *Lupinus* genus, as well as some legumes such as chickpea (*Cicer arietinum* L), red clover (*Trifolium pratense* L), and faba beans (*Vicia faba* L) can enhance P uptake from different soil P-pools by enhancing soil exploration. Hansen et al. (2022) reported that radish (*Raphanus sativus* L.) had higher P-uptake efficiency than other plant species tested in soils with a high abundance of Fe and Al-bound P. Nevertheless, the specific preference and uptake efficiency of this crop for P species bound to either Al or Fe in acidic soil has not been widely investigated.

Therefore, the present study aims to investigate and compare the effectiveness of a cover crop (radish) and a cash crop (maize) in reclaiming legacy P. By using synchrotron-based X-ray fluorescence spectroscopy mapping and X-ray absorption near edge structure speciation analysis, we evaluated the amount, and the chemical species of legacy P depleted and absorbed by these crops to understand whether it is P-bound with Fe or Al. We hypothesized that (1) Synchrotron X-ray fluorescence mapping has the potential to reveal distinct elemental profiles within the rhizosphere of plant; (2) the cover crops known for effectively reclaiming legacy P will demonstrate unique patterns of P distribution and chemical composition in the rhizosphere, unlike less efficient crop species.

## MATERIAL AND METHODS

### Isothermal adsorption of P onto Fe and Al minerals

Hematite and boehmite were selected as representatives of Fe and Al oxide and oxyhydroxide minerals respectively in soil (Prietzel et al. 2016). Hematite was purchased from Sigma Aldrich while boehmite was obtained from Sasol. They were used as P adsorbents and the P adsorbed on them was used as the only source of P in the substrate prepared, mimicking the legacy P in soil.

For the adsorption, 12.8g of each mineral sample was weighed and mixed in an Erlenmeyer flask with 800 mL 0.01 M KCl solution containing 5 g L^-1^ P. The P solution was prepared from a mixture of 21g (4.78 g L^-1^ P) of potassium dihydrogen phosphate (KH_2_PO_4_) and 1.24g (0.22 g L^-1^ P) of dipotassium hydrogen phosphate (K_2_HPO_4_) and kept at pH between 5.5 to 6. The P concentration (5 g L^-1^) was chosen to attain maximum and complete sorption of the mineral’s surface according to the adsorption capacities obtained in a preliminary sorption test. After mixing, the suspensions were incubated at 70°C with vigorous 2 minutes shaking once a day for 20 days, as an adaptation of the procedure described by Barrow et al. (2022). Thereafter, the supernatant solutions were first decanted to reduce the liquid volume. Subsequently, the sedimented minerals were rinsed with deionized water to remove all weakly adsorbed P from the minerals, the solids were separated by centrifugation at 2320 g for 10 minutes. The P-loaded minerals were frozen at -20°C for 48h and then freeze-dried. The quantity of P sorbed onto the minerals was determined colorimetrically after digesting the sample in HNO_3_, as described by Wieczorek et al. (2022).

### Preparation of P-containing substrate for rhizosphere assessment

We prepared a substrate consisting of 50% coarse sand (0.5mm to 1mm), 25% kaolinite (purchased from Sigma Aldrich), 9% hematite-P, and 16% boehmite-P. This mixture proportion was used to simulate the composition of a 50% clay Oxisol (Darunsontaya et al. 2010), as well as to mimic the presence of legacy P into the soil. The final substrate contained a total of 1480 mg P g^-1^, being 50% bound to hematite and 50% bound to boehmite. The sand used was initially analyzed for total P and found to contain 45 mg P kg^-1^. To remove this P, the sand was soaked in 0.1M HCl for 48h, thereafter washed with deionized water and oven-dried (35°C, 24h). The washing procedure stopped when the concentration of P leached by HCl was below 1.43 mg P kg^-1^ which corresponds to the detection limit of the ammonium molybdate spectrometric method (Santos de Souza et al. 2023). This step was essential to ensure that the Fe/Al-P complexes were the only sources of P for the plants, allowing us to accurately study the uptake of the P adsorbed onto the minerals. Sand was also sterilized to eliminate any microorganisms’ activities.

### Experiment 1: Rhizosphere study

#### Experimental Design and Planting Technique

The rhizosphere of the radish and maize was investigated by cultivating them in rhizoboxes (265mm x 250 mm x 25mm, height, width, and depth, as described in Figure 1A). Three treatments were established: (i) maize plant, (ii) radish plant, and (iii) control (no plant). Four independent replicates for each treatment were employed.

**Figure 1.**
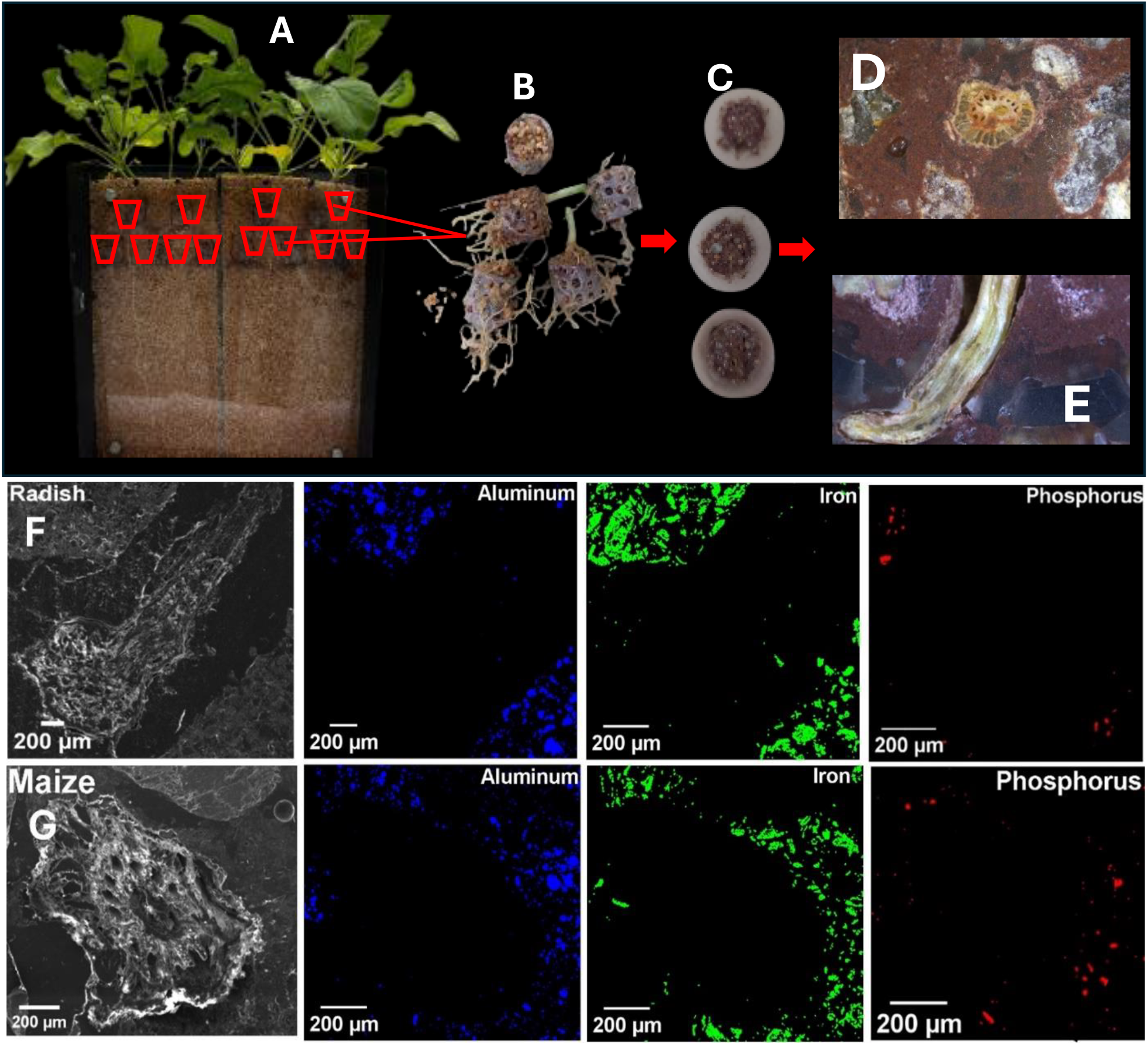
Experiment 1 set-up and sample preparation: Radish grown in rhizobox device (A). Each rhizobox has dimensions 265mm x 250mm x 25mm (height x width x depth) with a transparent removable plastic of 3mm thick. Each rhizobox was divided into two separate compartments, with a middle plastic rod of 5mm thickness separating them. Each of these compartments was then considered a replicate. To investigate the rhizosphere of the crops, we made use of a small cylinder of height 13mm and a diameter of 15mm. These cylinders were perforated and filled with the prepared substrate and strategically positioned within the rhizoboxes beneath the seed during planting to enable root penetration. The remaining space in the rhizobox was filled with sand. The cylinders with the roots after disassembling the rhizobox (B). Sliced cylinders contain root (C). Petrographic-like thin sections contain maize root (D), and radish root (E) viewed with stereo microscope. Scanning electron micrographs images and corresponding energy dispersive X-ray spectroscopy (EDS) elemental distribution maps of radish (F) and maize (G) roots showing aluminum (blue), iron (green), and phosphorus (red) in the rhizosphere. The maps illustrate the distribution of the elements in the rhizosphere of radish and maize.

To investigate the rhizosphere of these crops, the substrate described above was loaded into perforated plastic cylinders 13mm in height and 15mm in diameter (Figure 1B). The cylinder containing the P-bearing “Oxisol-like substrate” acted as a “root trap”. They were placed in the rhizobox to allow the penetration and establishment of roots as well as the mass flow of nutrients. The remaining space in the rhizobox was filled with sand.

Two seeds were planted at 2-3cm below the soil surface and just above the cylinder filled with P-rich substrate. The rhizoboxes were placed in a plant growth chamber under controlled conditions for 5 weeks (14 h illumination, 27°C, 70% relative air humidity). They were kept at an angle of 45° to stimulate roots to grow downward. Water was supplied from the top and the amount was calculated from the sand water holding capacity and the gravimetric water content in the rhizobox was maintained at 80% of the water holding capacity by weighing. Irrigation was done with a nutrient solution once a day during the first week, increased to twice a day after the second week due to the very low water-holding capacity of the sand. All nutrients, except P, were supplied by the nutrient solution of concentration; Ca(NO_3_)_2_.4H_2_O (230g/L), KCl (7g/L), H_3_BO_3_ (0.6g/L), MgSO_4_ (26g/L), K_2_SO_4_ (90g/L) MnSO_4_.H_2_O (0.17g/L), CuSO_4_.5H_2_O (0.025g/L), ZnSO_4_.7H_2_O (0.288g/L), NH_4_NO_3_ (28.8g/L), (NH_4_)_6_Mo_7_O_24_.4H_2_O (0.012g/L) and Fe-EDTA (33.2g/L), while the only source of P was the substrate in the perforated cylinders. This experiment was carried out using 4 biological replicates, each replicate consisted of two plants, and for each plant, we employed 3 root traps substrate-containing cylinders (Figure 1A).

#### Sample preparation for rhizosphere assessment

The experiment was terminated in the 5^th^ week after planting, the rhizoboxes were disassembled and the cylinders containing roots (Figure 1B) were carefully removed to avoid disturbing the roots within. For morphological and spectroscopy analysis, the sample preparation was done following the standard method used in geological thin sectioning (Camuti and Mcguire 1999). Five cylinders per treatment were selected, dried at room temperature, and impregnated under vacuum with epoxy resin Araldite 2020/A and 2020/B, then stored under the vacuum at room temperature for 24 hours to remove bubbles and air from the system. After hardening, the cups were sliced transversally (Figure 1C) using a slicing machine (rectilame 1 23 03). The samples were then cut with a diamond saw to have a petrographic-like thin section of approximately 1.5mm thick.

#### Chemical composition of the rhizosphere

Separately, some root-trapped cylinders containing roots were selected, the substrate attached to the roots was carefully collected, and the rhizosphere substrate was then ground to powder, and a FlashSmart Elemental Analyzer was used to determine total carbon exudated by the plants. Fourier-transform infrared (FT-IR) spectra were also recorded on the finely ground rhizosphere substrate with an alpha II FT-IR spectrometer within a spectral range of 400 – 4000 cm^-1^ and a spectral resolution of 2 cm^-1^ to determine the chemical composition of the rhizosphere.

#### Morphological and Spectroscopic Characterization

The samples (petrographic-like thin section) were first observed under a stereo microscope (Wild M8), representative samples are shown in Figure S1 of the electronic supplementary information (ESI). Then, high-resolution images were acquired using the scanning electron microscope (SEM), the samples were coated with carbon, and the images were recorded under 10 keV. Energy-dispersive X-ray spectroscopy (EDS) was also recorded at 25 keV.

X-ray fluorescence microanalysis (µXRF) measurements of the rhizosphere samples were conducted at the Tarumã end-station of the Carnauba beamline at the Brazilian Synchrotron Light Laboratory (LNLS, Brazil) and the ID- 21 beamline of European Synchrotron Radiation Facility (ESRF, France). In this later, X-ray absorption near edge structure (µXANES) was also explored. Both µXRF and µXANES were recorded at, nearby, and far away (500 µm) from the roots, as illustrated in Figure S2. All analyses were carried out using 3 replicates per treatment.

At the Carnaúba beamline, the µXRF maps were recorded using a 0.2 × 0.5 µm monochromatic X-ray beam at 7.2 keV focused on the sample surface using a Kirkpatrick–Baez (KB) mirror system. The two-dimensional 20 × 250 µm chemical maps were recorded in flyscan mode, at a 1 µm pixel resolution. The XRF spectra were recorded by two four-element silicon drift detectors (Vortex-ME4, Hitachi High-Technologies Science America, USA) in a He- doped atmosphere to increase the sensitivity to light elements.

At the ID21 beamline, the measurements were carried out using a 0.35 × 0.7 µm monochromatic X-ray beam operating at 2370 eV and focused by KB mirrors. Especially herein, the third harmonics at 7.2 keV were not rejected to allow the simultaneous detection of Fe, as well as other elements with smaller z. The analyses were performed under a vacuum to minimize scattering and absorption by air. The XRF maps were acquired by measuring 120 × 120 µm areas with a 2 µm lateral resolution and a 200 ms pixel^-1^ dwell time. The µ-XANES spectra were collected within a 2.141 - 2.21 keV energy range, at 0.2 eV energy step.

#### Data processing

Data from µXRF was processed using PyMCA software (Solé et al. 2007). Elemental fluorescence intensity peak signals were fitted in PyMCA to obtain net elemental distribution maps that were then overlayed as RGB image maps. ImageJ software was used to apply a two-dimensional Euclidean distance to the XRF intensity map containing part of the root to measure the distance from the root to the pixels in the substrate neighboring the root. Four distance classes were defined: 0-10µm from the root, 10-20µm, 20-200µm and 500-740µm from the root. The image statistic tool in PyMca was used to determine the average elemental count rate at the defined spots, these qualities are directly proportional to the elemental concentration. The µXANES spectra acquired from three distance classes were processed using Athena with the Demeter 0.9.26 package (Newville and Ravel 2005). Background subtraction and normalization were performed, then principal component analysis (PCA) was applied. Linear combination fitting (LCF) of P-XANES data was performed reference standards to examine P speciation close to the root and far from the root using Athena software over an energy range of 2140 to 2200 eV. Inorganic phosphate sorbed to Fe oxide (hematite), Al oxyhydroxide (boehmite), and kaolinite standards were measured and used to align the recorded XANES spectra with the beamline’s reference compounds library employed for linear combination fitting (LCF) analyses.

### Experiment 2: Phosphorus extraction study in tubes

#### Experiment setup

This experiment aimed at determining the fraction of P absorbed by maize and radish from hematite-P and boehmite-P. The plants were grown in 50 mL polycarbonate tubes containing a 50g mixture of the substrate (Figure S3). The composition of the substrate and the solution employed in irrigation is presented in Table 1. Tubes were kept at 80% water-holding capacity during the experiment and placed in a growth chamber in the same condition as the rhizoboxes described above. The experiment was carried out using five biological replicates, i.e. five (5) tubes per crop species for a certain substrate composition.

**Table 1:**
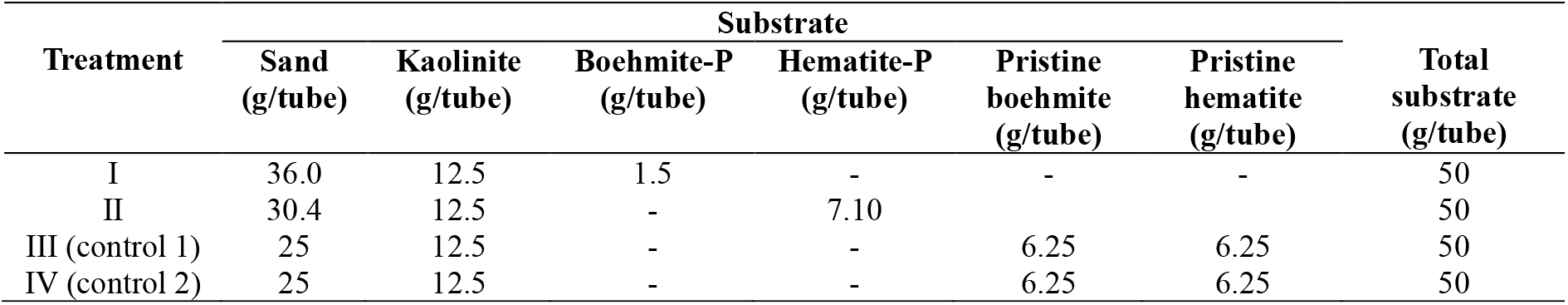
Summary of the tube experiment treatments. For treatment I and II, the amount of mineral-P complex (hematite-P and boehmite-P) added to the mixture of sand and kaolinite was calculated to supply 25 mg of P while the amount of sand was calculated to reach a final mass of 50g. Treatments I, II, III (control 1) were irrigated with a nutrient solution without P. While treatment IV (control 2) was irrigated with a complete nutrient solution with P.

| Treatment | Substrate |  |  |  |  |  | Total substrate (g/tube) |
| --- | --- | --- | --- | --- | --- | --- | --- |
|  | Sand (g/tube) | Kaolinite (g/tube) | Boehmite-P (g/tube) | Hematite-P (g/tube) | Pristine boehmite (g/tube) | Pristine hematite (g/tube) |  |
| I | 36.0 | 12.5 | 1.5 | - | - | - | 50 |
| II | 30.4 | 12.5 | - | 7.10 | - | - | 50 |
| III (control 1) | 25 | 12.5 | - | - | 6.25 | 6.25 | 50 |
| IV (control 2) | 25 | 12.5 | - | - | 6.25 | 6.25 | 50 |

#### Plant sampling and analysis

The plants’ older leaves were pruned biweekly to enhance mineral-P extraction. The experiment was terminated, and the plants were harvested 90 days after planting. At harvest, the shoots and roots were separated, and roots were carefully rinsed with deionized water, then dried at 70°C for 72 h and weighed to quantify the biomass production, then ground for analysis. Phosphorus concentrations in shoot and root tissues were determined colorimetrically after total digestion using HNO_3_ (Wieczorek et al. 2022). To calculate the amount of P taken up by the plant from the P strongly adsorbed onto the minerals, the total P obtained from the control plants that did not receive any P application was subtracted from the total P in the plant tissue planted on the mineral-P complexes, while the percentage of P recovered was calculated using the following equation (Amadou et al. 2022):

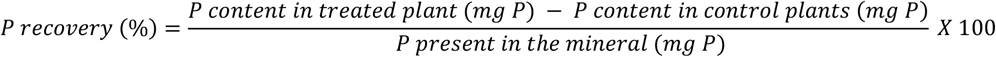

#### Statistical analysis

The quantitative data such as data on plant biomass, plant tissue P content, P uptake, and total carbon exudated by the plant root were subjected to one-way analysis of variance (ANOVA) using R software (R Core Team 2019). Duncan’s multiple range test was conducted to compare the treatment means that showed significant differences. Differences between means were considered significant when *p* < 0.05. All data visualization, including graphs and bar charts was performed using OriginPro, version 2024b (OriginLab Corporation, Northampton, MA, USA)

## Results

### Experiment 1: Elemental distribution in the rhizosphere

The scanning electron micrographs (SEM) images and corresponding energy dispersive X-ray spectroscopy (EDS) elemental distribution maps recorded at the root-substrate interface of radish and maize plants are shown in Figures 1F and 1G, respectively. It is worth noting that the void between the root and the substrate which appears as a gap immediately after the root surface is due to shrinkage of the root after drying prior to resin infiltration. This is because the root tissue contracts during the drying process, and when resin is applied, it fills the resulting space around the root. The substrate immediately beyond the resin (where the element signals for Al, and Fe are detected) corresponds to the first contact with the root in its natural state. Thus, the element distribution observed in the EDS maps (blue for Al, green for Fe, and red for P) reflects the initial interactions between the root and the surrounding substrate.

The µXRF maps presented in Figures 2 and S4-S5 details the distribution of Al, Fe, and P at the interface and further from the root and substrate for both radish (Figure 2A) and maize (Figure 2B). Figures 2C and 2E present the maps for radish and maize, respectively, covering sections of the root, adjacent substrate, and regions further from the root. The root segment appears as the P-rich region in the top left corner of Figure 2C (radish) and Figure 2E (maize). Figures 2D and 2F display maps recorded 500 µm away from the root for radish and maize, respectively, revealing a highly heterogeneous P distribution.

**Figure 2.**
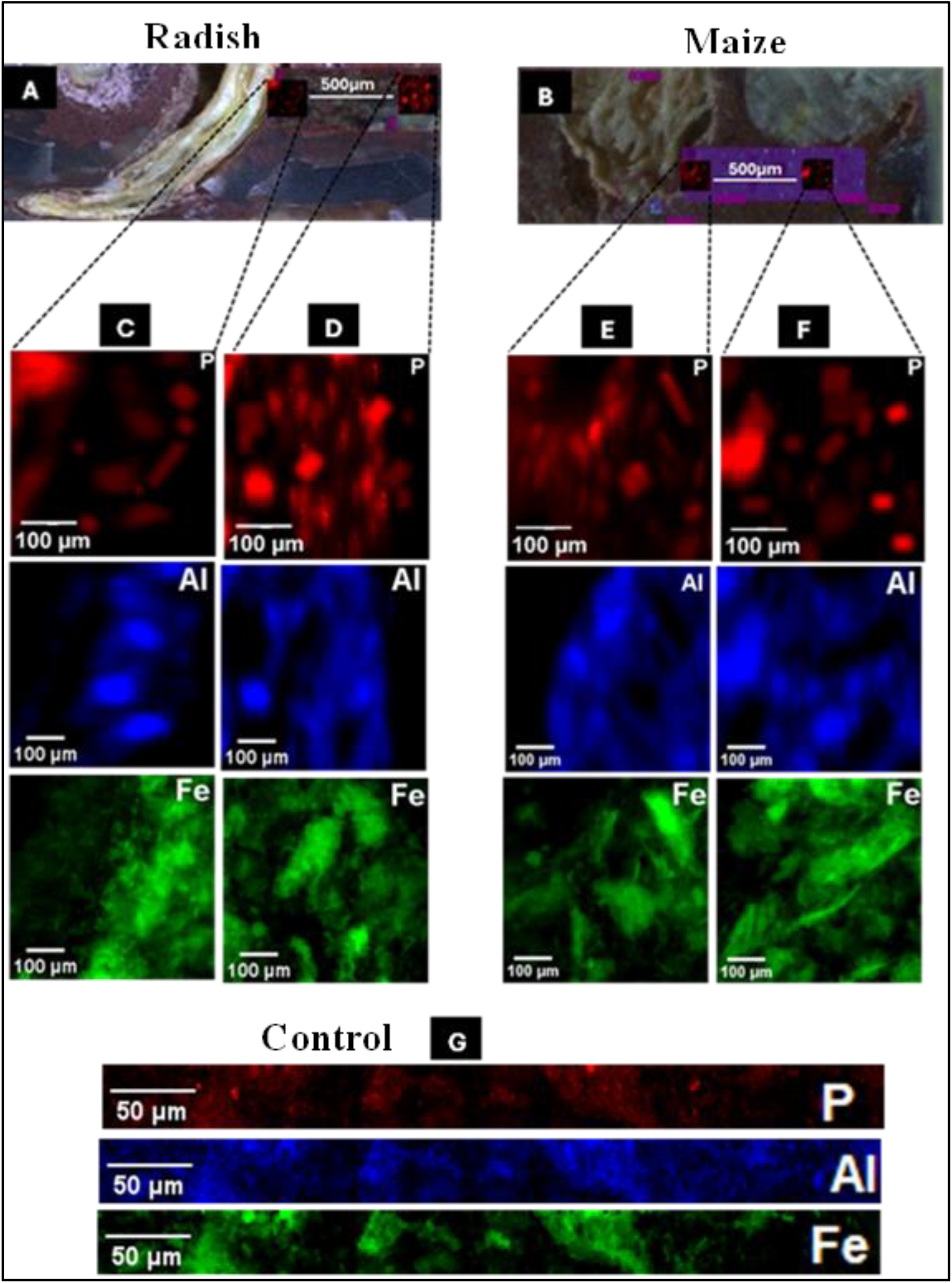
Photographs and micro X-ray fluorescence (μ-XRF) distribution maps of Aluminum (blue), Iron (green) and Phosphorus (red) at the rhizosphere of radish and maize plants and map of control (without plant). A and B are the photographs showing the root of radish and maize respectively and large map of size 58 × 200µm^2^, step size 7 × 7µm and 0.1second dwell time within which two other high resolution small maps each of size 120 × 120µm^2^, step size 2µm were taken. For radish and maize, maps C and E include part of the root section and neighboring soils (substrate), from where three distinct distances range from the root were defined namely, 0-10µm, 10-20µm, and 20-200µm distance away from the root section. The root segment represents the P-enriched region seen at the top left of map C and E. Radish map D and maize map F were recorded at a distance 500µm away from the plants root. Map G is the map of control (20 × 250 µm^2^, and step size 1µm).

A notable difference in P distribution is evident between radish and maize maps, both close to the root and far away from the root. For radish plants, the P distribution is lower near the root compared to areas further away, suggesting the existence of depletion zones and active P uptake by the plant. The P distribution near the root overlaps with regions of high Fe concentration, indicating the co-localization of Fe and P close to the root. Some P spots are also visible, localized far from the root and overlapping with both Fe and Al. However, the co- localization of P with Al is not evident in the immediate vicinity of the root but becomes more prominent moving away from the root. In contrast, Figure 2G shows the control map with a uniform distribution of P alongside Fe and Al, highlighting the absence of plant influence on P distribution.

#### P mobilization in the rhizosphere of radish and maize

Figure 3 shows the X-ray fluorescence counts of P associated with Al and Fe across the defined distances from the root of radish, maize, and the control (substrate without plant). The bars indicate that there is a progressive increase in the concentration of P associated with Al with distance from the root in both radish and maize. For radish, the P counts associated with Al (Figure 3 A) increases from 315 counts at 0-10 µm up to more than 1200 counts at 500- 740 µm from the root. Also, for maize (Figure 3 C), there is an increasing trend, from 600 at 0-10 µm to more than 1500 counts at 500 – 740 µm from the root. Radish plants have a lower overall P content associated with Al at the root zone compared to maize, suggesting some differences regarding the Al–P uptake ability of the two crops. A similar trend is observed with P counts associated with Fe, however, in radish (Figure 3 B), P on Fe increases from 500 at 0-10 µm to about 1300 counts at 500-740 µm from the root. Maize (Figure 3 D) appears to still follow such a trend but has recorded slightly higher P concentration which gradually increased to almost 1400 at the furthest distance recorded from the root. For the control (Figure 3 E, F), there is a uniform distribution in the concentration of P associated with both Al and Fe across all distances. The count of P is roughly constant around 8-9 × 10^4^ with no significant changes.

**Figure 3.**
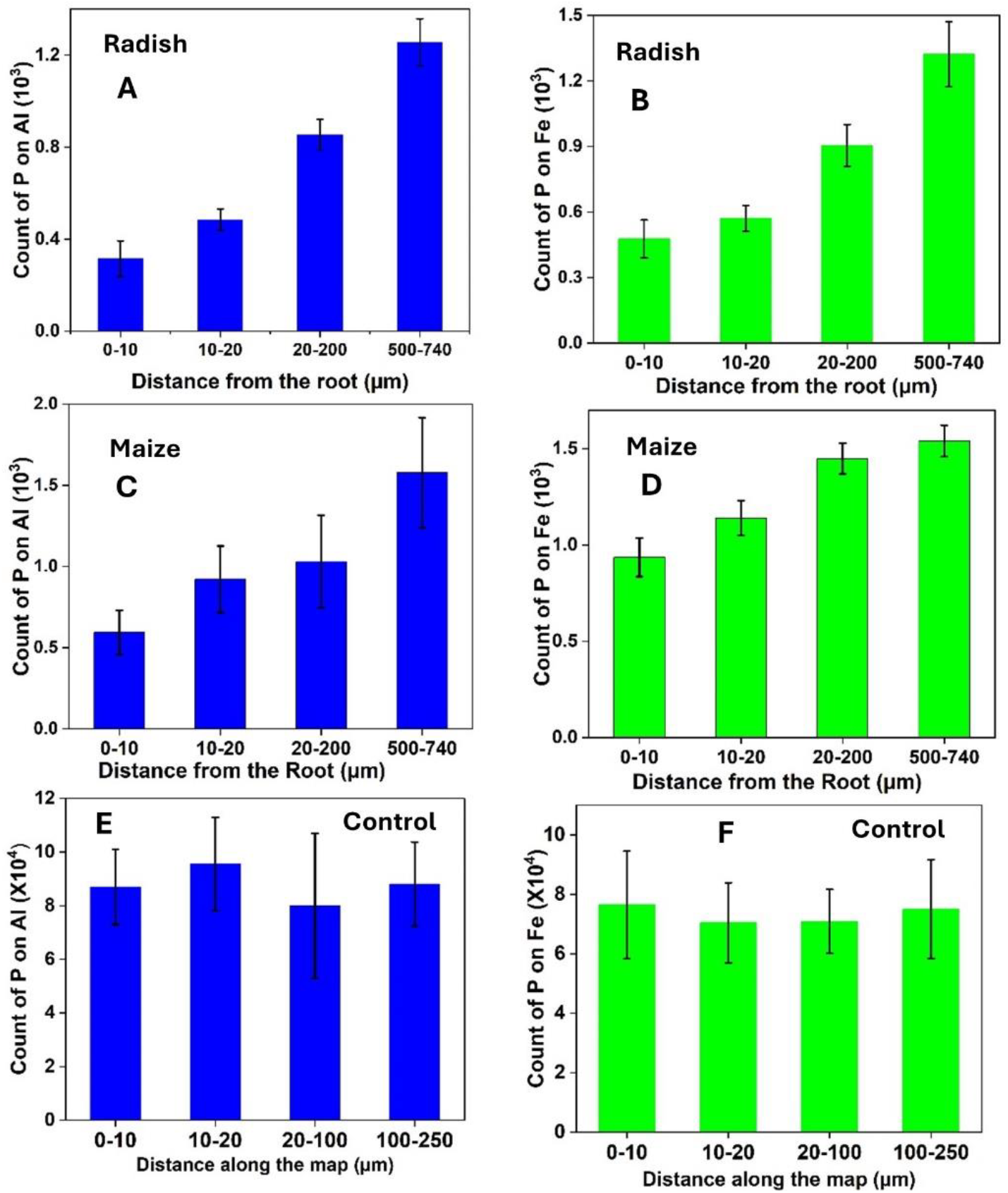
Phosphorus XRF pixel counts associated with aluminum (Al) and iron (Fe) at distinct distances from the root of radish (A and B) and maize (C and D), in comparison with the control (E and F). Radish and maize show a decrease in counts of Al- and Fe-associated P with distance to the root, indicating P uptake by the plant. Radish exhibits the lowest P concentration associated with Al close to root compared to maize which exhibits higher overall P counts, suggesting greater efficiency in accessing P from Al and Fe complexes. In the control, P count distribution remains uniform across all distances, indicating the immobility of Al- and Fe-bound P in the absence of plant roots. The bars represent the average values with the standard deviation.

#### Phosphorus K-edge XANES

Representative samples of P XRF maps of radish and maize with selected points of interest (POI) for µ-XANES are presented in Figures 4AB-5AB. The principal component analysis was applied on all the normalized P K- XANES spectra acquired to evaluate if there is any variation among the XANES spectra acquired at the root-soil continuum and at a distance far away from the root. The PCA scatter plots in Figure 4C and 5C show eigenvalues obtained from the PCA of the spectra collected at distinct distance from the root as shown in Figure 4A, 4B for radish and 5A, 5B for maize. For radish (Figure 4C), principal component 1 (PC1) explains 88.3% of the total variance in the spectra, and PC2 explains only 5.3%. While for maize (Figure 5C), PC1 also accounts for 90.7% of the total variation of all the spectra, and PC2 accounts for 3.7%. K-mean clustering was applied to the data to quantitatively group the data points to have cluster of similar spectra. Interestingly, three different clusters were obtained represented with different colors (Figure 4C and 5C). First cluster shown in green, second in red, and third in blue which were the spectra acquired at 0-20µm, 20-200µm, and 500-740µm distance from the roots, respectively.

**Figure 4.**
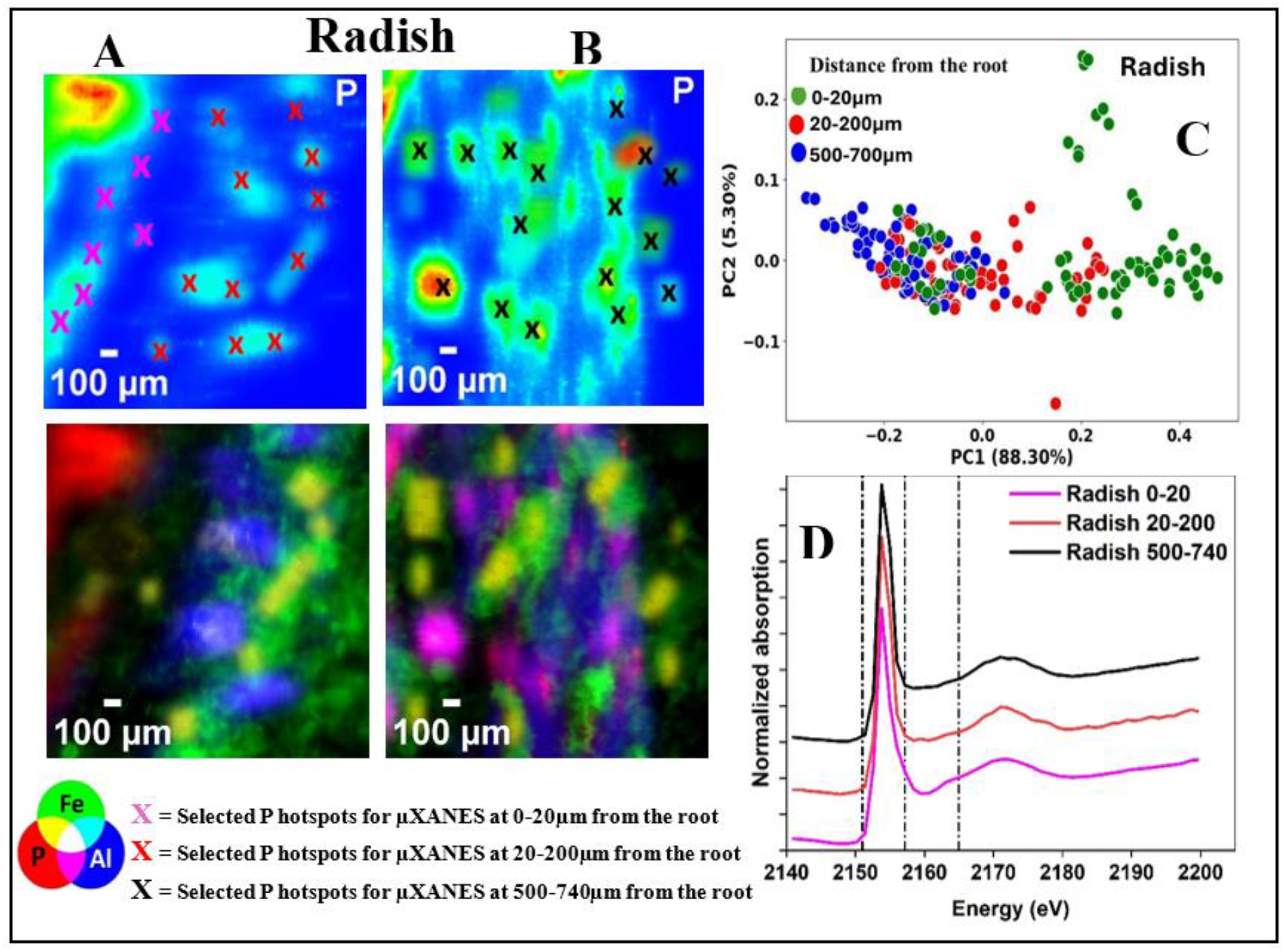
XRF maps (A and B; 120 × 120 µm^2^, 2 µm step) of P with selected points of interest for µ-XANES for radish. Map A include part of the root (the P-rich area at the top left), map B was recorded at 500µm away from the root. Principal component analysis (PCA) and µXANES scatter plot of PC1 vs. PC2 from the radish samples µXANES spectra. The color shows batches differences of cluster for P species at different distances from the root (C). Normalized P K-edge XANES spectra (points) of each cluster group from the PCA analysis (D). The RGB images show the co-localization of Al-P-Fe

**Figure 5.**
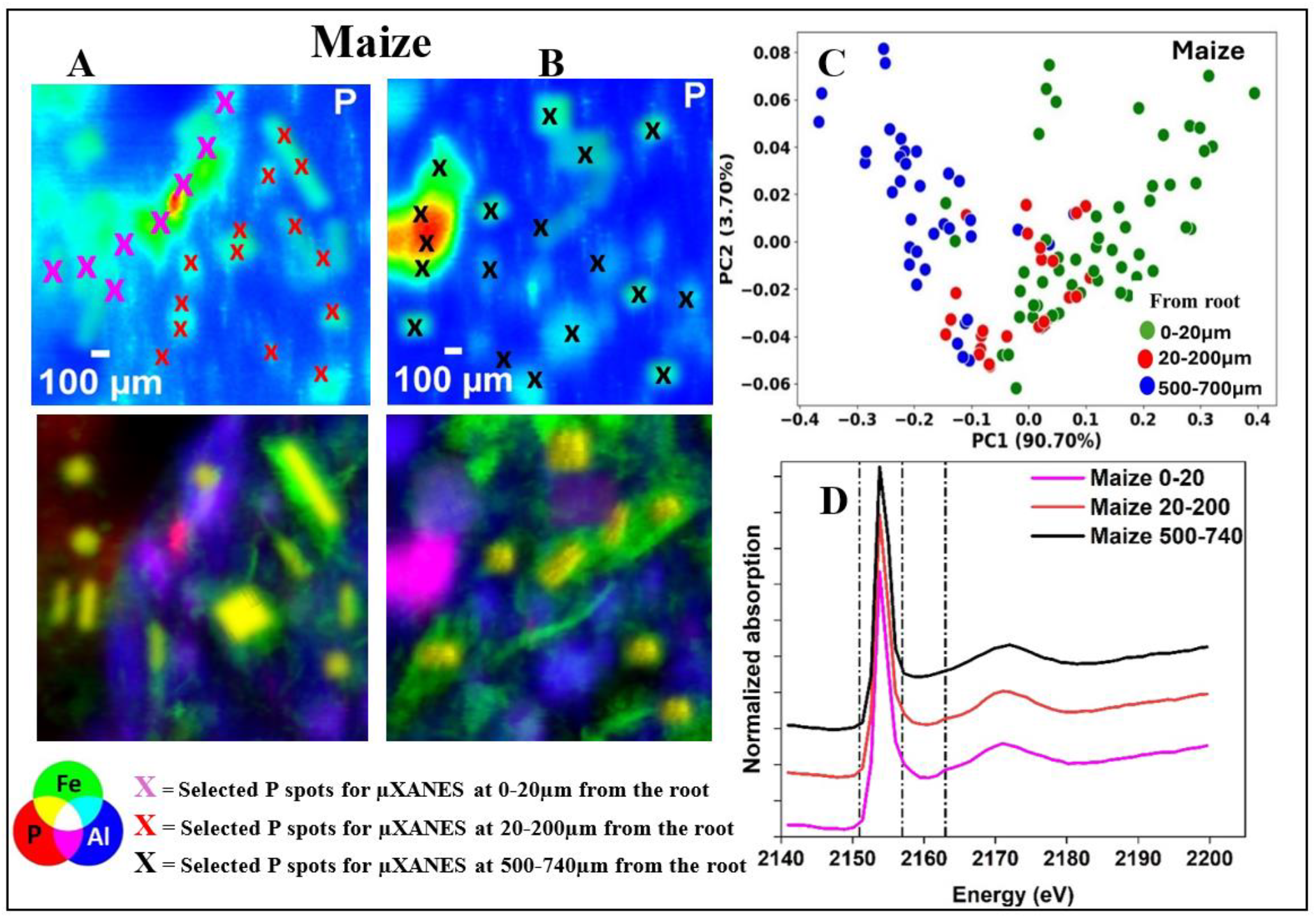
XRF maps (120 × 120 µm^2^, 2 µm step) of P with selected points of interest for µ-XANES for maize. Map A include part of the root (the P-rich area at the top left), map B was recorded at 500µm away from the root. Principal component analysis (PCA) and µXANES scatter plot of PC1 vs. PC2 from the maize samples µXANES spectra. The color shows batches differences of cluster for phosphorus species at different distances from the root (C). Normalized P K-edge XANES spectra (points) of each cluster group from the PCA analysis (D). The RGB images show the co-localization of Al-P-Fe

#### Linear combination fitting (LCF)

The spectra of the standards compound used in linear combination fitting (LCF) of the P K-XANES clusters obtained from the PCA analysis are available in the supplementary material (Figure S6). The bulk P K-edge XANES spectra for the three clusters for radish and maize are provided in Fig 4D and 5D, respectively. A pre-edge peak feature between 2148 and 2152 eV and weak post-edge shoulder, that are strong indication of P sorbed on Fe as presented in the reference spectra (Figure S6) were observed in the XANES spectra acquired at 0-20µm distance from the root of both radish and maize (Figure 4D and 5D). However, this pre-edge peak feature becomes weaker with increasing distance from the root, as noticed with spectra acquired at 20-200µm away from the root for the two crops. Spectra recorded at 500-700µm away from the root of both radish and maize have no pre-edge peak which is a feature of P adsorbed on Al oxyhydroxide and a small post-edge shoulder which is feature of P associated with Fe. The curve fitting for each cluster obtained from the PCA analysis are provided in supplementary material in Figure S7 and Table S1. The LCF results showed that those P spots recorded at 0-20µm from the root of both radish and maize are predominated with P sorbed on Fe-Oxide (hematite-P), accounting for about 79% for radish and 67% for maize (Figure S7A, S7B). However, the results also provided that the P sorbed on Al oxyhydroxide (boehmite-P) standard accounted for 56% P at 20-200µm from the root of radish with small fraction of P adsorbed on kaolinite (Figure S7C). In contrast, P sorbed on Fe-oxide standard was found to dominate 20-200µm from the root of maize and accounted for 70% P followed by P sorbed on boehmite (28%) and small proportion on kaolinite (Figure S7D). There were no visible spectra differences between the 500-740 µm distances from the roots of radish and maize, as both contained similar proportions of the standard compounds (Figures S7E, S7F). At this distance, P adsorbed on Al oxyhydroxide was the dominant form (51% for radish and 54% for maize), followed by P adsorbed on Fe-oxide (46% for radish and 43% for maize), with a very small proportion of P adsorbed on kaolinite (3% for both radish and maize).

#### Chemical composition of the rhizosphere

The infrared spectra recorded on the rhizosphere shows a distinct chemical composition of the substrate (Figure 6 A). As expected, due to the composition of the substrate and as extensively reported in literature, the absorbance bands observed at 3680-3651 cm^-1^ correspond to the external OH groups in the Al-OH, and Si-OH groups that take part in forming bond between octahedral and tetrahedral sheet layers in the kaolinite clay (Rouibah et al. 2024). Peaks at 3280, 3087, and 1075 cm^-1^ are three strong absorption bands that characterize pure boehmite. The two former peaks are assigned to the asymmetric and symmetric O-H stretching vibrations from (O)Al-OH while the later bands are attributed to symmetric bending of Al-OH (Nkoh et al. 2022). The strong band at 793 cm^-1^ observed in control which become weak and shifted 726 cm^-1^ in maize and radish could be attributed to stretching vibrations of Fe-O (Yang et al. 2023). Interestingly, a new absorbance signal peak at 1628 cm^-1^ was observed in the rhizosphere of radish. Figure 6B presents the total carbon exudated by the radish, and maize. The highest total carbon content was obtained in the rhizosphere of radish (0.15%), which is significantly higher, about 2.14-fold than that of maize (0.07%).

**Figure 6.**
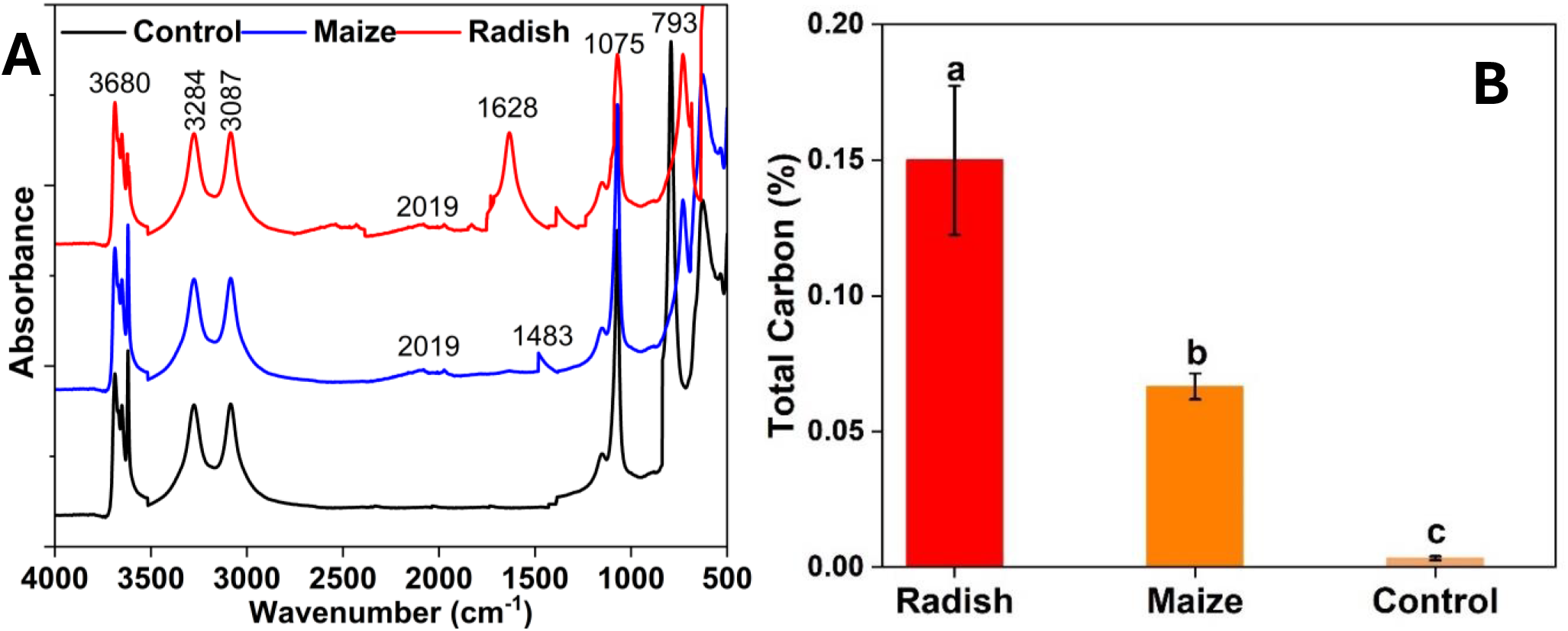
Fourier transform infrared spectra of the rhizosphere of radish, maize and the soil from the control (A). Total carbon exudated by the plants (B).

### Experiment 2: Phosphorus extraction study

#### Biomass Production

Table 2 summarizes the effect of different treatments on total biomass (shoot + root biomass) production by radish and maize. For all the treatments, shoot biomass was always significantly higher than root biomass in both crops. The highest total biomass production was obtained for both radish and maize when they were grown in control II, where the P source was soluble, from a complete nutrient solution, this shows that a full nutrient balance is crucial for maximum biomass production. The lowest total biomass was found when both crops were grown in control I, where no P was applied, with the absence of P greatly limiting plant growth. Radish grown in the substrate containing P sorbed on boehmite produced more total biomass (2.57±0.23g) than P sorbed on hematite (1.80±0.28g), which account for 42.78% increase in total biomass. This difference indicates that P from boehmite may be more readily available compared to P from hematite. The maize biomass production was almost equal or slightly higher when grown in the substrate containing P sorbed on boehmite (5.58±0.56g) compared to when grown in the substrate containing P hematite (5.37±0.57g), with just a 3.91% increase in total biomass production.

**Table 2:** Effect of different Treatments on shoot, root, and total biomass of radish and maize. The results represent mean ± standard deviation. Means with a different letter were significantly different according to Duncans Multiple range test at P < 0.05. Comparisons of means were presented for each plant with treatment category.

| Treatment | Radish |  |  | Maize |  |  |
| --- | --- | --- | --- | --- | --- | --- |
|  | Shoot biomass (g) | Root biomass (g) | Total biomass (g) | Shoot biomass (g) | Root biomass (g) | Total biomass (g) |
| I | 2.43±0.24b | 0.14±0.08b | 2.57±0.23b | 4.84±0.54b | 0.74±0.09b | 5.58±0.56b |
| II | 1.72±0.26c | 0.08±0.03c | 1.80±0.28c | 4.46±0.59c | 0.91±0.23c | 5.37±0.57c |
| III (control 1) | 0.28±0.18d | 0.08±0.06d | 0.36±0.22d | 0.49±0.11d | 0.16±0.07d | 0.65±0.18d |
| IV (control II) | 4.01±0.46a | 1.10±0.15a | 5.11±0.21a | 6.34±0.38a | 2.63±0.13a | 8.97±0.09a |

Treatments with the same letter within a parameter are not significantly different, while different letters signify significant differences.

#### Total P uptake and recovery from the P adsorbed minerals

Total P uptake and percentage P recovered by both radish and maize grown under various treatments are presented in Figure 7. In the bar graph (Figure 7A), as expected, radish grown in substrate irrigated with a complete nutrient solution (control II) showed the highest P uptake (12 mg P), which was significantly greater than other treatments. Maize grown under the same conditions takes 8 mg P, even lower than radish planted in P adsorbed on boehmite.

**Figure 7.**
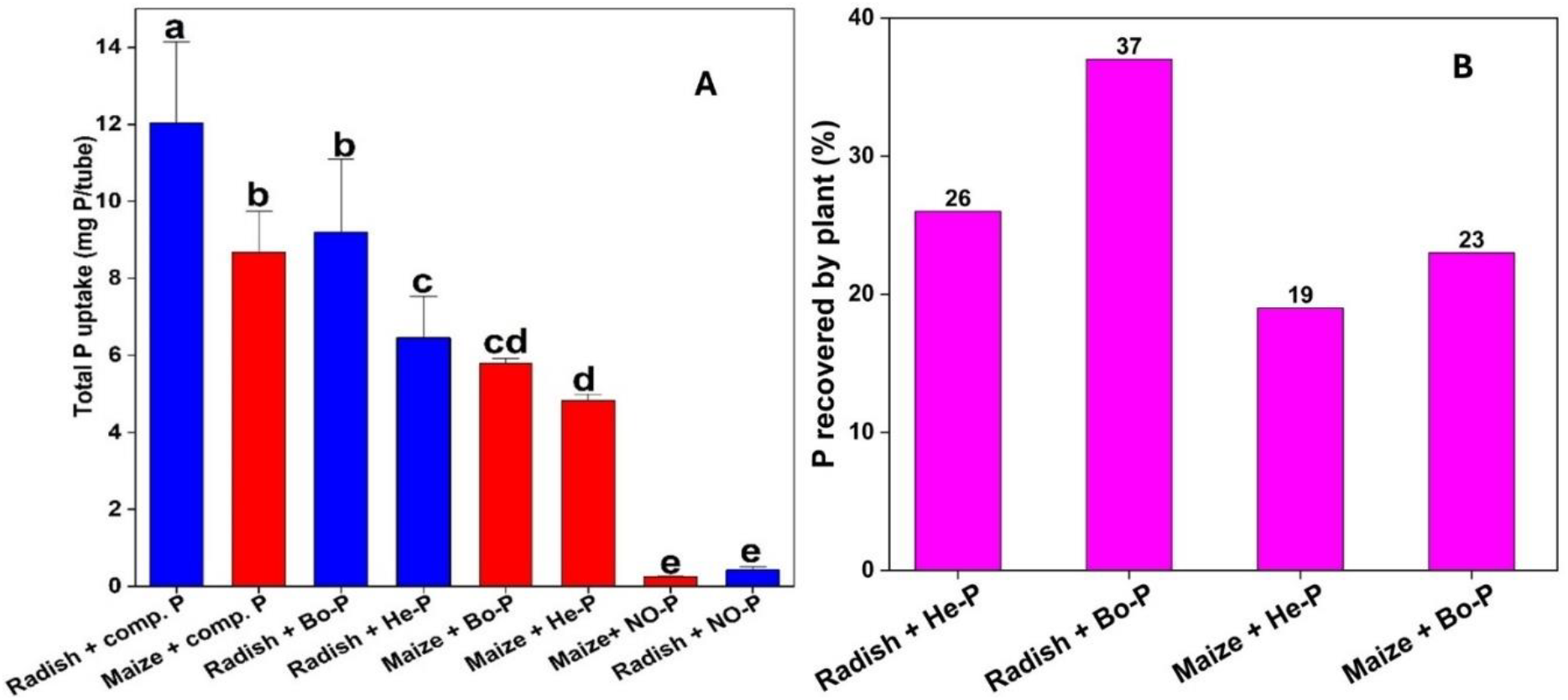
Total P uptake by maize and radish per tube (A). Percentage of total P recovered by maize and radish (B). Radish +Bo-P and Maize +Bo-P are radish and maize plants respectively, planted in substrate contains mixture of pure sand, strongly adsorbed P on boehmite (25mg P), kaolinite and irrigated with nutrient solution without P. Radish + He-P and Maize + He-P are radish and maize plants respectively, planted in substrate contains mixture of pure sand, strongly adsorbed P on hematite (25mg P), kaolinite and irrigated with nutrient solution without P. Radish + NO-P and Maize + NO-P (Control 1) are radish and maize plant respectively planted in a substrate contains mixture of pure sand, pure hematite, pure boehmite, and pure kaolinite irrigated with nutrient solution without P. Radish + comp. P and Maize + comp. P (Control 2) are radish and maize plant respectively, planted in a substrate contains mixture of pure sand, pure hematite, pure boehmite, and pure kaolinite irrigated with complete nutrient solution with P. The error bars indicated the standard deviation of replicates (n=5). Distinct letters on the bars indicate significant differences between treatments (P < 0.05).

For P adsorbed minerals, radish extracted significantly more P from boehmite-P, with a total uptake of 9.19 mg P per tube (Figure 7A), compared to 6.46 mg from hematite-P (P<0.05). In contrast, maize showed no significant difference in total P uptake between the boehmite-P and hematite-P treatments (P<0.05). Notably, total P uptake by radish from boehmite-P was twice what maize uptake from hematite-P. However, there was no significant difference between the P uptake by radish from hematite-P and that uptake by maize from boehmite-P.

Radish demonstrated a notably higher P recovery efficiency, recovering 37% of the P adsorbed onto boehmite and 26% of the P adsorbed onto hematite, compared to maize, which recovered only 23% and 19% for the same treatments, respectively (Figure 7 B). Overall, radish demonstrated significantly higher total P uptake and recovery efficiency than maize, with the greatest difference in boehmite-P treatment.

Phosphorus concentrations in the shoots of radish were significantly higher than those in the shoots of maize in both boehmite-P and hematite-P substrates. However, there were no significant differences in the P concentrations within the roots of radish and maize for either treatment (Figure S8).

## Discussion

The results obtained from the distribution of Al, Fe, and P in the rhizosphere of both plants (radish and maize) confirmed our first hypothesis that synchrotron X-ray fluorescence mapping has the potential to reveal distinct elemental profiles within the plant’s rhizosphere. The distribution of chemical maps of Al, Fe, and P exhibited heterogeneity in the distribution of these elements, with observed variations in P concentration near and far from the roots for both species, which is consistent with other spectroscopic investigations of elemental distribution at the root-soil interface (Muccifora et al. 2021; Van Veelen et al. 2020)

The P X-ray fluorescence counts is directly proportional to the concentration of P. In our study, the average P counts close to the root were significantly lower compared to far away from the root. This shows the establishment of a P depletion zone and P uptake by the plants because a visible concentration gradient is observed which is a driving force for diffusion (Marschner and Rengel, 2011). We observed that the P depletion layer is not restricted to soil solution, but also involving P sorbed onto the minerals. Moreover, the P depletion in the root zone was more pronounced in radish than in maize, suggesting a higher P uptake in radish compared to maize plants. This aligns with previous studies highlighting the ability of radish to take up more P from less labile P fractions when compared to other crops (Soltangheisi et al. 2020). The ability of radish to outperform maize in P uptake can be attributed to specific and pronounced chemical interactions that occurred within the rhizosphere, primarily driven by root exudation processes, such as exudation of organic acids that are very efficient in overcoming the P binding mechanisms (Jones 1998). Studies have shown that P mobilization involves several strategies which include organic acid exudation by plant root (Almeida et al. 2020; Pantigoso et al. 2020).

Our results show that radish exuded organic acids, as evidenced by the FTIR spectra. The absorption band observed at 1628 cm^-1^ in the radish rhizosphere soil, which was not detected for maize, could be attributed to the vibration of C=O of carboxylic acid, which has been reported to occur at 1600cm^-1^ to 1630cm^-1^ wavenumber (Chua et al. 2020; Gebauer et al. 2018; Guo et al. 2021). Furthermore, the ability of radish to exudate a larger amount of total carbon compared to maize could support the evidence of organic acid production as it was estimated that total carbon contains about 5 to 10% organic acids (Calvo and Vallejo, 2002; Pantigoso et al. 2020). Higher P uptake by radish when compared with other crops was also observed by Moreira de Carvalho et al. (2008) from Brazilian Oxisols attributed to the ability of the crop to exude malic acid. In a similar study, Pavinato et al. (2008) evaluated plant extracts of six plant species including radish and maize, in which radish released the highest amount of malic and citric acid, referring to its efficiency in enhancing soil P availability. Therefore, it could be explained that the organic acid exuded by radish compete with P adsorbed on the surface of hematite and boehmite or complex the Fe and Al ions to enhance P desorption (Jones 1998; Pavinato and Rosolem, 2008) thereby increasing the concentration or availability of P in the solution (Rao et al. 2016). Additionally, the acidic environment created by acid exudation could lower the rhizosphere pH, further enhancing the dissolution of Al-P and Fe-P complexes. The ability of maize to uptake some P might be associated with its ability to release some carbon exudates (Van Veelen et al. 2020).

Our results indicate that P bound to Al is significantly depleted by radish, with a narrow concentration distribution near the root compared to areas farther from the root, creating a concentration gradient. This suggests that radish likely depletes more of Al-bound P for its demand, as opposed to Fe-bound P, which shows a shallower concentration and distribution gradient. In contrast, near the maize root, the concentration of Al-bound P is higher compared to what is observed in radish, indicating that maize also extracts Al-bound P but appears to be less dependent than radish. It is clearly shown that the plant roots influence the localization and release of P linked to Al and Fe. Both radish and maize can extract it, although radish does this more efficiently over a longer distance. The uniform distribution of P in control suggests that P associated with Al and Fe is immobile in the absence of roots, proving how effective rhizospheric processes are in mobilizing P. These results are consistent with the previous studies which show that roots alter the chemical P environment in their vicinity (Bakker et al. 2013; George et al. 2024; Saleem et al. 2018).

The PCA and the LCF further support the results obtained from elemental mapping. The distinctions observed between the P K-XANES spectra, concerning the distance from the root indicate that the plant’s activity has a major effect on the P speciation within the rhizosphere (van Veelen et al. 2020). For radish, the LCF revealed that most of the P close to the root was P sorbed to Fe with a small amount of P sorbed to Al and no trace of P sorbed onto kaolinite, while in maize, P adsorbed on Fe was also dominant with a higher percentage of P sorbed on Al compare to what was found in radish, and some amount of P adsorbed on kaolinite. This suggests that both Fe-P and Al-P were mobilized by the plants but there is a higher uptake of the Al-P compared to Fe-P.

Interestingly, the biomass production and P uptake data from the second experiment also corroborate the above-identified differences in the P acquisition strategies between radish and maize. Radish grown in substrate containing Al-P produced more biomass (43% increase) than in Fe-P. This means that radish more efficiently utilizes the Al-P nutrient, and as it were, is consistent with µXRF mapping and LCF results depicting that radish depleted more of Al-P. There was, however, no significant difference in the biomass production of maize when grown in boehmite and hematite bound P suggesting that maize deficiency is more equilibrated or less specific in its P acquisition of Al-P and Fe-P. The total P uptake and P recovery results also confirm that radish can effectively extract P from Al-bound minerals than from Fe-bound P. Radish was able to extract 9.19 mg P from boehmite-P compared to hematite-P (6.46 mg P), while maize could not distinguish the two substrates in terms of P uptake. Moreover, radish also exhibited higher P recovery efficiency (37% from boehmite and 26% from hematite) compared to maize (23% and 19%, respectively). This shows that radish is better suited for recovering P from strongly bound forms in soils dominated by Al oxyhydroxide, making it a potentially valuable crop in P-deficient soils with high Al content.

The higher uptake of P adsorbed onto Al compared to Fe may be attributed to the specific organic acids released by radish, which preferentially solubilize Al-bound P. Zhang et al. (1997) identified tartaric, malic, and succinic acids as the dominant organic acids produced by radish, particularly under P deficient conditions, these acids were found to enhance the mobilization of Al-P, as radish utilized P from aluminum phosphate (AlPO_4_) more efficiently than from calcium phosphate (Ca_3_(PO_4_)_2_) in a quartz sand culture. In contrast, Zhang et al. (1997) also reported that rapeseed (*Brassica napus*) predominantly released citric and malic acids, which facilitated greater solubilization of calcium-bound P. He linked these differences in P solubilization to the specific organic acids exuded by each crop, underscoring their role in determining the plant’s P source preferences. Another possible reason could be due to the nature of the surface complexes formed. Organic substances released by the plants may solubilize Al-bound phosphate due to the weaker binding energy of monodentate or bidentate-mononuclear Al-P surface complexes (Kim and Kirkpatrick 2004), which makes it more soluble than Fe-P (Shane et al. 2008) as opposed to the stronger bidentate-binuclear surface complexes typically formed between P and Fe (Kwon & Kubicki, 2004). These stronger Fe-P interactions make Fe-bound phosphate less susceptible to solubilization, resulting in a lower uptake by the plants. This is in accordance with Adam (2016), who added citrate to the mixture of P sorbed Fe oxide mineral (ferrihydrite) and P sorbed boehmite minerals and found that phosphate was preferentially released from boehmite mineral, concluding that it might be related to a stable configuration of P on the iron mineral.

Furthermore, previous spectroscopic findings of phosphate sorption reported that phosphate bound with Fe oxide at pH between 4 and 6 form stable bidentate-binuclear surface complex (Khare et al. 2007; Luengo et al. 2006), while bidentate mononuclear surface complex formation was reported for phosphate sorbed on Al (Yoon and Bleam 2009). Parfitt (1989) reported that the binding energy follows the order: monodentate > bidentate > binuclear complexes, while the likelihood of phosphate desorption increases in the opposite order. Far away from the root, at 500-740 µm distances from the root, both radish and maize exhibit similar P speciation characterized by Al and Fe forms showing that minimal plant effect on the proportion. Thus, the specific mineralogical composition of the soil is likely to contribute to the effectiveness of P recovery, with certain crops (such as radish) more suited to soils rich in Al-bound P. In this regard, understanding these can guide the selection of crops. The ability of radish to mobilize legacy P could have significant implications for enhancing PUE in agriculture particularly in soils with high Al-bound P, while contributing to sustainable soil fertility management. The use of radish could also ensure a good cycling of nutrients, mitigating the reliance on synthetic P fertilizers to reduce the environmental impact of agricultural systems.

Synchrotron-based µXRF mapping and µXANES analysis applied in this present work offered us much relevant information about P distribution and speciation in the rhizosphere. The techniques provide unprecedented spatial resolution and elemental sensitivity which enable us to visualize the microscale distribution and detailed interactions between P, Al, and Fe in the rhizosphere of the studied plants (Hesterberg 2010). This enhances our ability to detect the P depletion zones and to interpret the plant-induced alterations in P availability. With µXANES analysis, we gained important insights into the species of P, e.g. Al-bound vs Fe-bound mostly uptake by our test plants. However, all scientific methodologies have inherent challenges, and this applies to this present work as well. Preparing large samples such as root-soil (rhizosphere) samples for high-resolution imaging is one of the biggest challenges due to limitations such as shrinkage of roots after root drying (Brunet et al. 2023). Nevertheless, these techniques remain powerful tools for elucidating elemental distribution and P speciation in the rhizosphere. The ability to link spatial distribution with chemical speciation gives a unique window into how plants access these tightly bound forms, as shown in this study. Future advancements in synchrotron and increased access could further enhance their utility and make them more accessible for agricultural research and sustainable soil management practices.

Lastly, several conditions used in this present work, such as substrate preparation (quartz sand and P-bound minerals) and controlled environmental conditions, may limit the generalizability of these findings to soil conditions in situ due to its heterogeneity; mixtures of organic matter, microorganisms, and variable minerals. Soil and environmental factors may influence the dynamics of root exudates, P mobilizing mechanisms and overall plant performance. Although the results of our findings generate useful insights into the processes controlling P uptake, additional research under in-field conditions is needed to confirm these findings and evaluate the transferability of the knowledge obtained across soil types and cropping systems. This method would serve as a bridge between laboratory experiments and commercial agriculture. Future research could deeper into the biochemical pathways that allow radish to preferentially take up Al-bound P. It would be beneficial to gain insight into the fine root exudation quantities by a deeper investigation of the biosynthesis pathways of organic compounds in radish roots, with comparative exudation patterns and their potential interaction with soil minerals. Additionally, advanced molecular techniques, such as transcriptomics and proteomics could help to identify the particular genes and enzymes involved in the production of those organic acids under limited P conditions.

## Conclusions

Our study shows the potential of employing synchrotron-based techniques to reveal the profile of elemental distribution in the rhizosphere and the explanation of the difference observed in the efficacy of radish and maize in mobilizing P from tightly bound forms. Radish compared with maize shows high efficiency in recovering more of the Al-bound P than Fe-bound P, what makes it a propitious cover crop that could be grown in soils with high legacy P, particularly where Al-bound P is the predominant, improving P use efficiency and recycling more this nutrient in the production system. The superior ability of radish to reclaim the legacy P is related to its ability to modify the rhizosphere chemistry by exudating larger amounts of organic molecules compared to maize.

## Abbreviations

µXRF: Microprobe X-ray fluorescence
µXANES: Microprobe X-ray absorption near edge structure
LCF: Linear Combination fitting
PCA: Principal Component Analysis
FTIR: Fourier-transform infrared spectroscopy
PUE: Phosphorus use efficiency
SEM: Scanning electron microscope
EDS: Energy dispersive X-ray spectroscopy

## Acknowledgments

This research used facilities of the Brazilian Synchrotron Light Laboratory (LNLS), part of the Brazilian Center for Research in Energy and Materials (CNPEM), a private non-profit organization under the supervision of the Brazilian Ministry for Science, Technology, and Innovations (MCTI). **Carlos Perez**, Carnauba beamline staff is acknowledged for assistance during the experiments 20232656.

We also acknowledge the European Synchrotron Radiation Facility (ESRF) for provision of synchrotron radiation facilities under proposal 97916 and we would like to thank **Hiram Michel-Castillo & Clement Hole** for assistance and support in using beamline ID21.

## Author contributions

All authors contributed to the conception of the study. Y.M.Y, H.W.P.C, and P.S.P. designed the study. Yahaya executed the experiments. Material preparation, data collection, and analysis were performed by Y.M.Y, H.W.P.C, G.S.M, and K.B. The first draft of the manuscript was written by Y.Y.M. Revision and approval of the manuscript were performed by H.W.P.C. All authors read and approve the final manuscript.

## Declarations

## Graphical Abstract

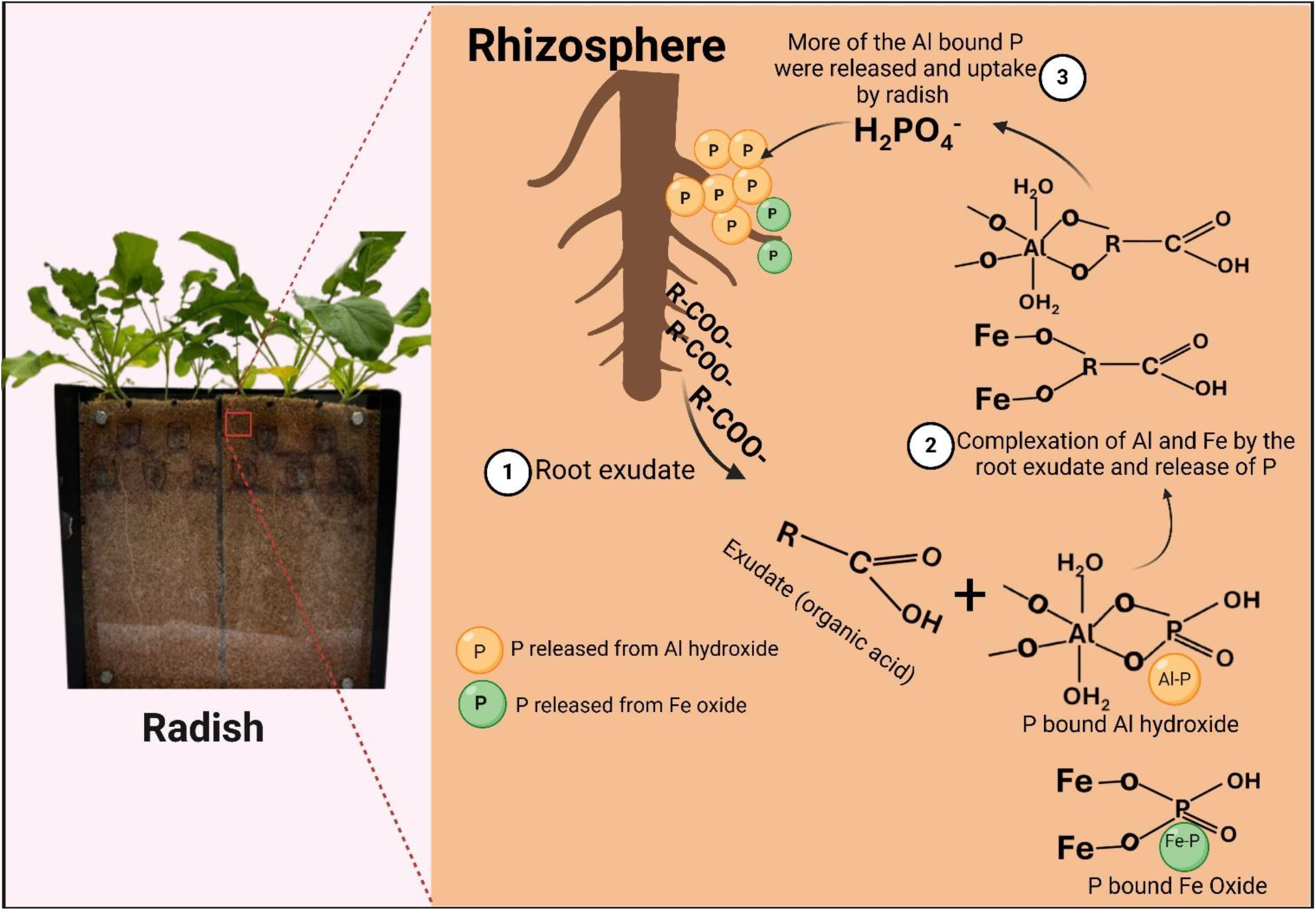

