## Supplementary figures for "How the rhizosphere chemistry explains the effectiveness of radish in reclaiming legacy phosphorus compared to maize"

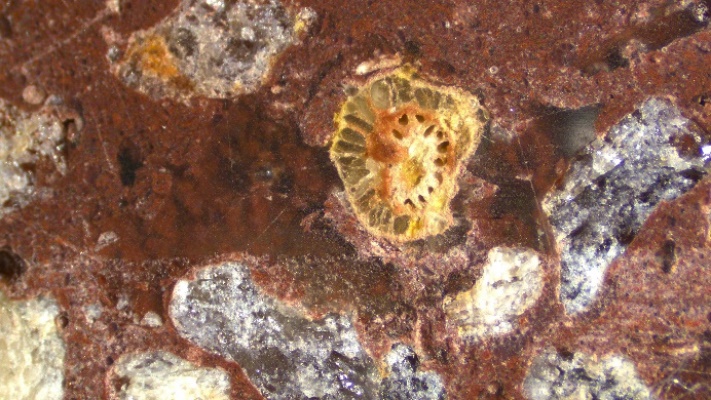

**B**

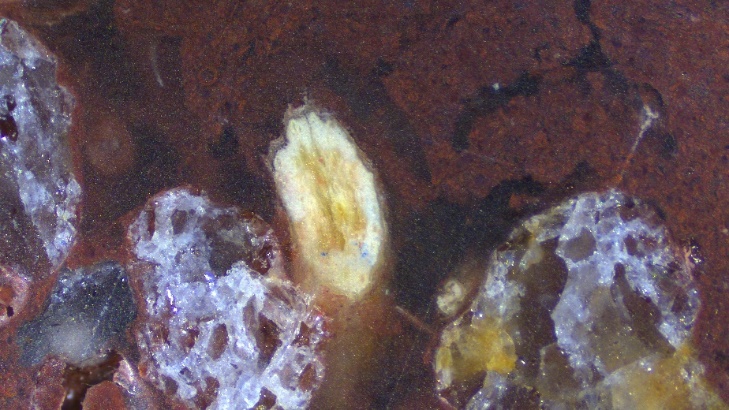

**A**

**Figure S1**. stereo microscopic images of prepared petrographic thin section showing (a) radish root (b) maize root

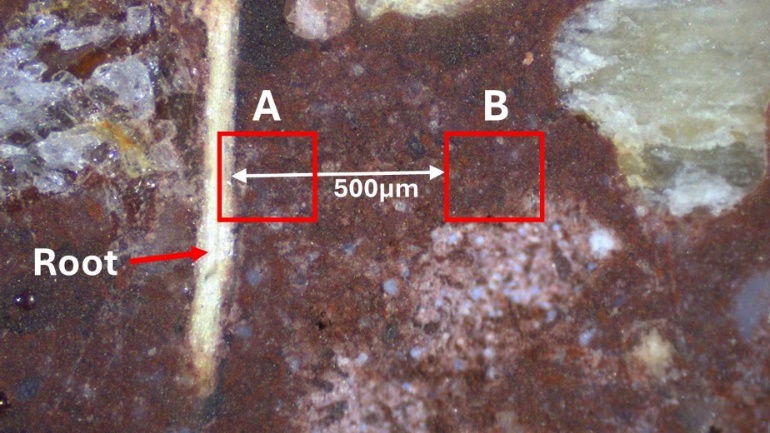

I

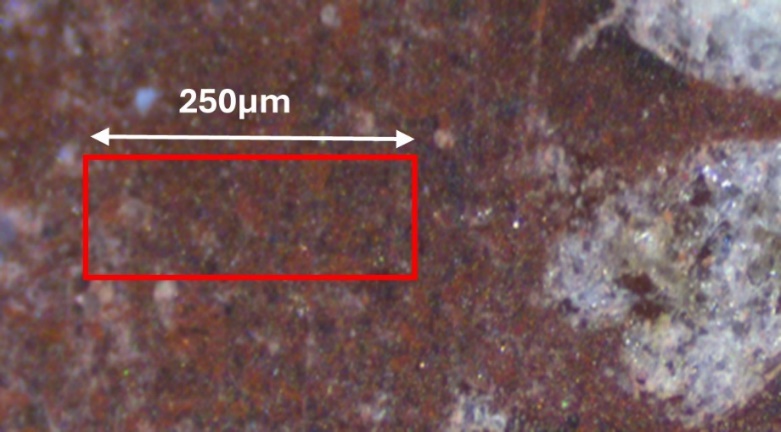

II

**Figure S2.** (I) Photograph illustrating the areas where high-resolution μXRF maps were recorded in the rhizosphere of radish and maize plants. Square A represents the area mapped with some part of the root surface, while square B illustrates an area mapped 500 µm away from the root. These regions were mapped and analyzed to compare the spatial distribution and concentration of phosphorus in proximity to the root against farther away from the root. (II) this illustrated how the control samples (sample without root) µXRF maps were recorded

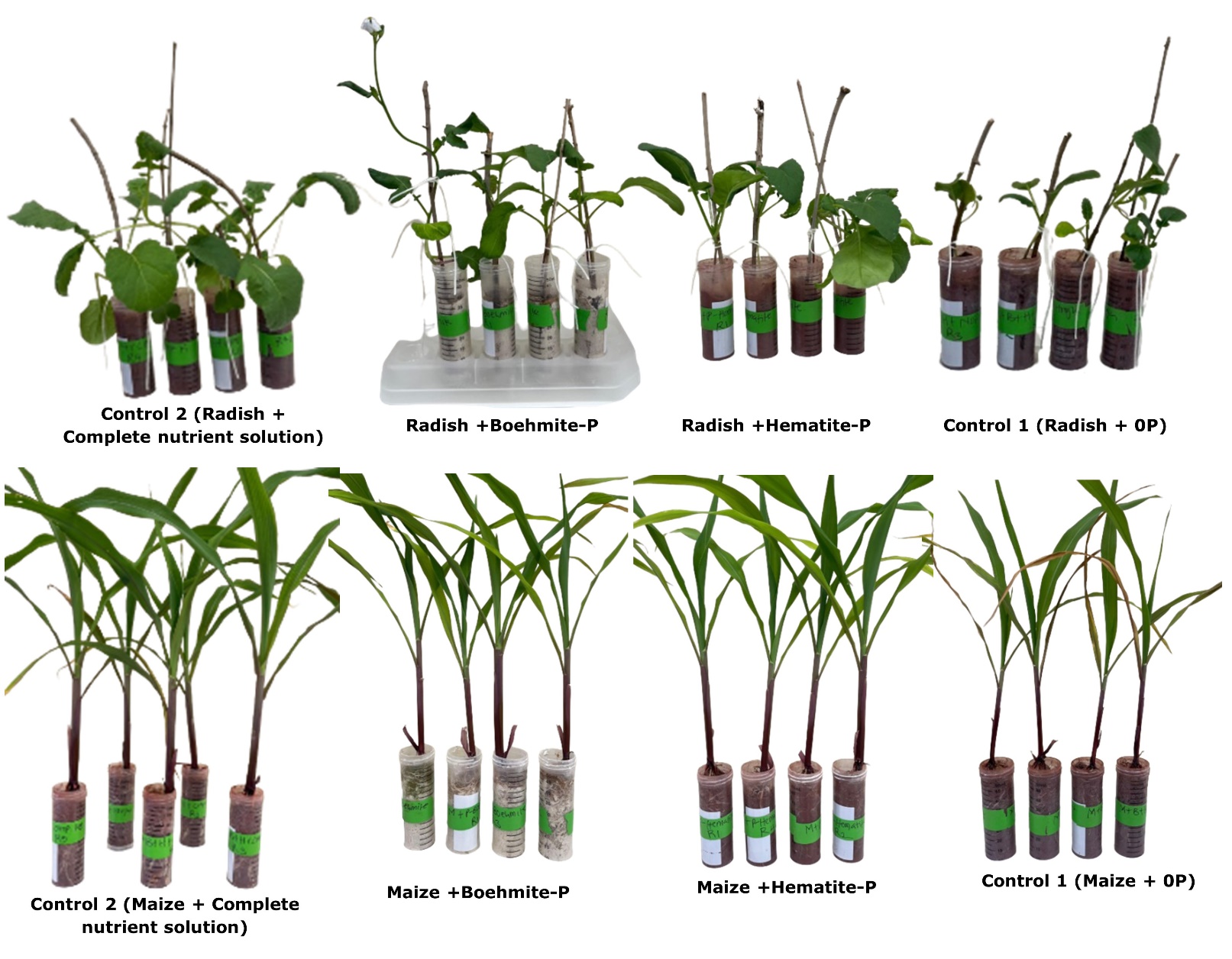

**Figure S3:** Experiment 2: tube experiment set up, extraction of P adsorbed on to minerals by radish and maize plant. Radish +Boehmite-P and Maize +Boehmite-P are radish and maize plants respectively, planted in substrate contains mixture of pure sand, strongly adsorbed P on boehmite (25mg P), kaolinite and irrigated with nutrient solution without P (treatment I). Radish +Hematite-P and Maize +Hematite-P are radish and maize plants respectively, planted in substrate contains mixture of pure sand, strongly adsorbed P on hematite (25mg P), kaolinite and irrigated with nutrient solution without P (treatment II). Control 1 (Radish +0P) and Control 1 (maize +0P) are radish and maize plant respectively, planted in a substrate contains mixture of pure sand, pure hematite, pure boehmite, and pure kaolinite irrigated with nutrient solution without P. Control 2 (Radish + complete nutrient solution) and Control 2 (maize + complete nutrient solution) are radish and maize plant respectively, planted in a substrate contains mixture of pure sand, pure hematite, pure boehmite, and pure kaolinite irrigated with complete nutrients solution with P

**C**

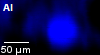

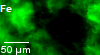

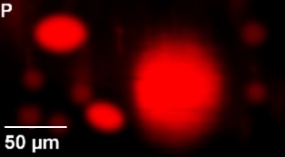

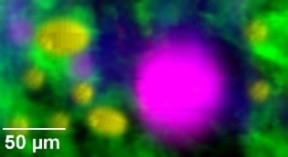

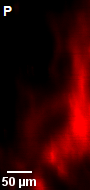

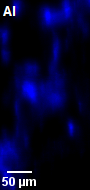

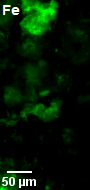

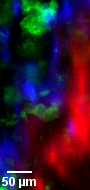

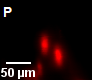

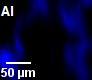

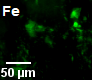

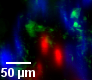

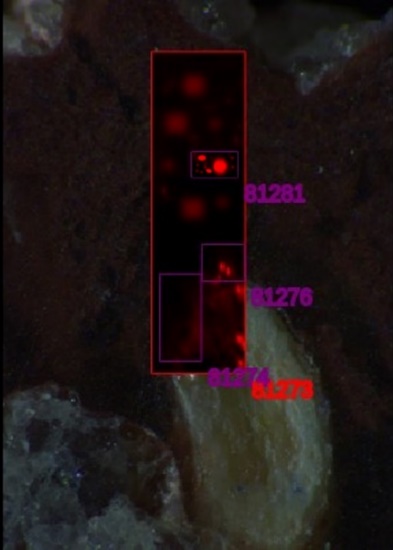

**B**

**D**

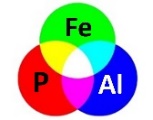

**A**

**Figure S4.** A replicate sample showing photographs and micro-X-ray fluorescence (μ-XRF) distribution maps of Aluminum (blue), Iron (green) and Phosphorus (red) at the rhizosphere of radish plant. A is the photograph showing the root of radish and a large map of size 58x200µm^2^, step size 7x7µm and 0.1second dwell time within which three other high resolution small maps B, C and D were taken. Maps B (size 90x190µm^2^, step size 2x2µm and 0.2 second dwell time) and map E (size 92x80µm^2^, step size 2x2µm and 0.2 second dwell time) include part of the root section and neighbouring soils (substrate), from where three distinct distances range from the root were defined namely, 0-10µm, 10-20µm, and 20-200µm distance away from the root section. The root segment represents the phosphorus-enriched region seen at the bottom right of map C and and center of map E. Map D (size 100x55µm^2^, step size 2x2µm and 0.2 second dwell time) was recorded at a distance 500µm away from the plants root. P depletion zone is clearly observed close to the root.

**B**

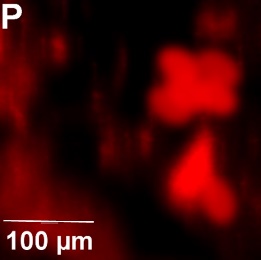

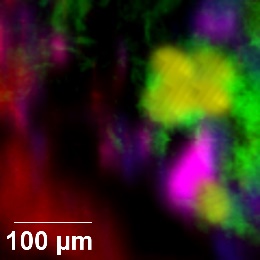

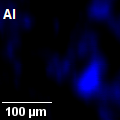

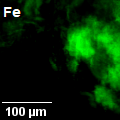

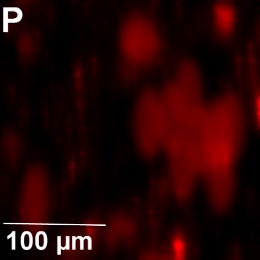

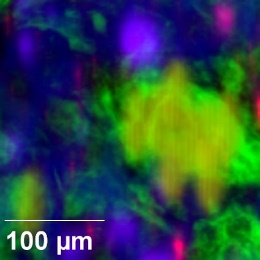

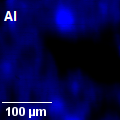

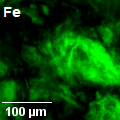

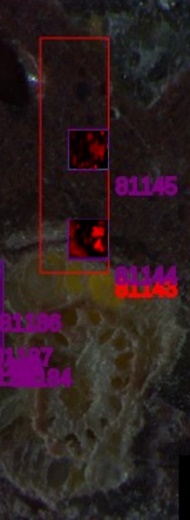

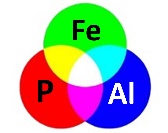

**A**

**C**

**Figure S5.** A replicate sample showing photographs and micro-X-ray fluorescence (μ-XRF) distribution maps of Aluminum (blue), Iron (green) and Phosphorus (red) at the rhizosphere of maize plant. A is the photograph showing the root of maize and a large map of size 58x200µm^2^, step size 7x7µm and 0.1second dwell time within which two other high resolution small maps B, and C (each with size 120x120µm^2^, step size 2x2µm and 0.2 second dwell time). Map B include part of the root section and neighbouring soils (substrate), from where three distinct distances range from the root were defined namely, 0-10µm, 10-20µm, and 20-200µm distance away from the root section. The root segment represents the phosphorus-enriched region seen at the bottom left of map B. Map C was recorded at a distance 500µm away from the plants root. P depletion zone is not clearly observed in maize as seen in radish.

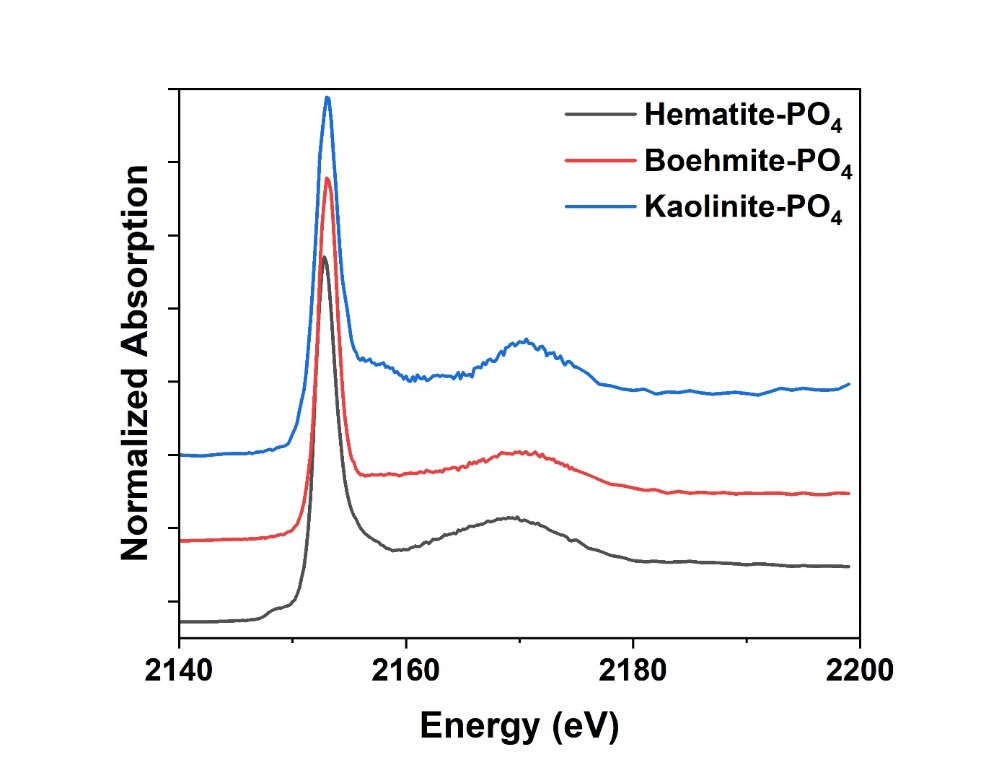

**Figure S6**. Normalized P K-edge XANES spectra of standard materials used for linear combination fitting of bulk XANES spectra. Hematite-PO_4_: phosphate-sorbed hematite, Boehmite-PO_4_: phosphate-sorbed boehmite, Kaolinite-PO_4_: phosphate-sorbed kaolinite

**Table S1.** Linear combination fitting (LCF) results as percentage of P adsorbed mineral and the uncertainties, with the R-factor, P adsorbed to hematite (Fe-P), P adsorbed to boehmite (Al-P), and P adsorbed Kaolinite (Kao-P).

|  |  | Phosphorus speciation (%) | | |  |
| --- | --- | --- | --- | --- | --- |
| Plant | Distance from the root (µm) | Fe-P | Al-P | Kao-P | R factor |
| Radish | 0-20 | 79±0.1 | 21±0.1 | 0.00 | 0.018 |
|  | 20-200 | 32±0.1 | 56±0.1 | 12±0.1 | 0.0026 |
|  | 500-740 | 46±0.1 | 51±0.1 | 3±0.1 | 0.0032 |
| Maize | 0-20 | 67±0.1 | 31±0.1 | 2±0.1 | 0.0047 |
|  | 20-200 | 70±0.1 | 28±0.1 | 2±0.1 | 0.0028 |
|  | 500-740 | 43±0.1 | 54±0.1 | 3±0.1 | 0.0027 |

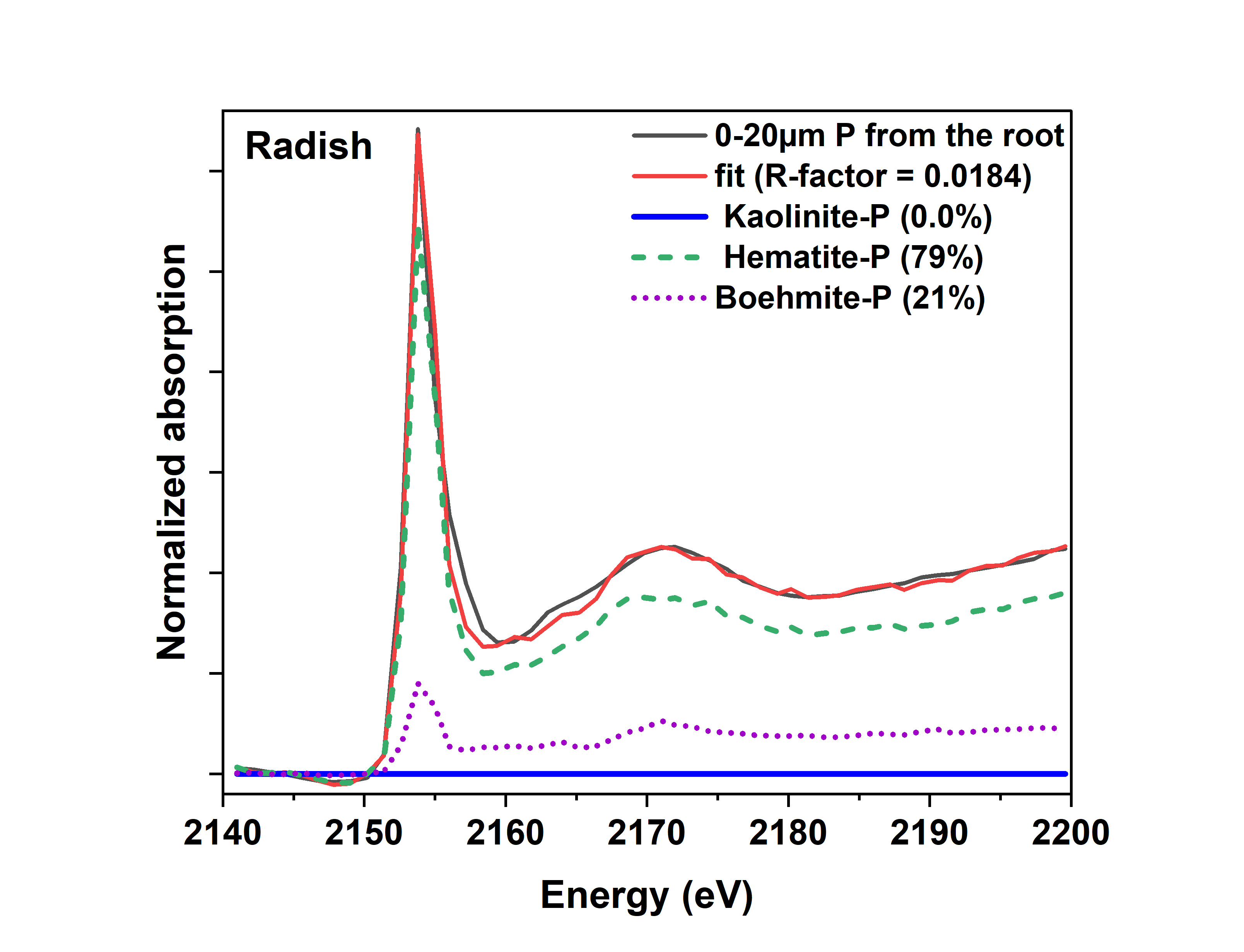

A

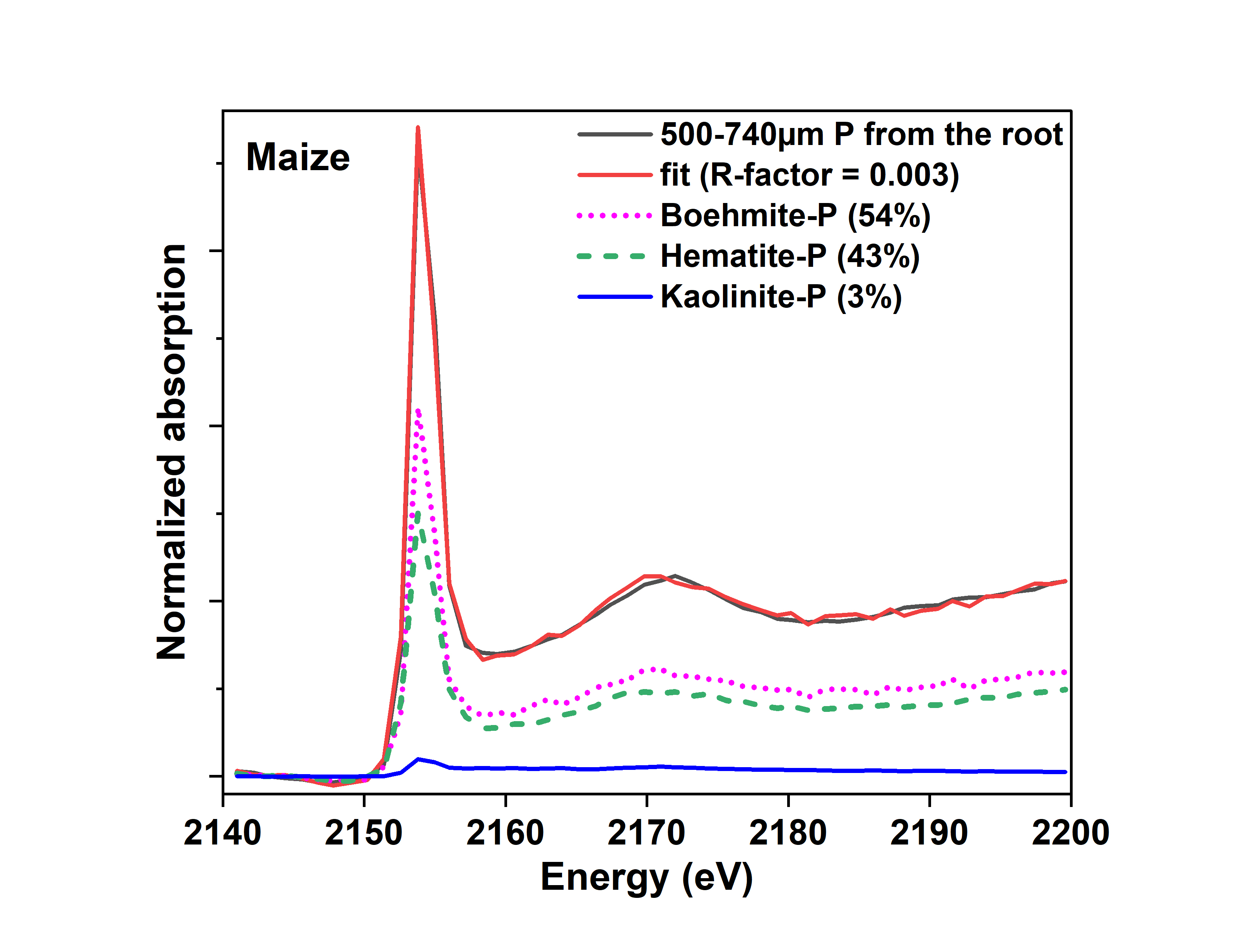

F

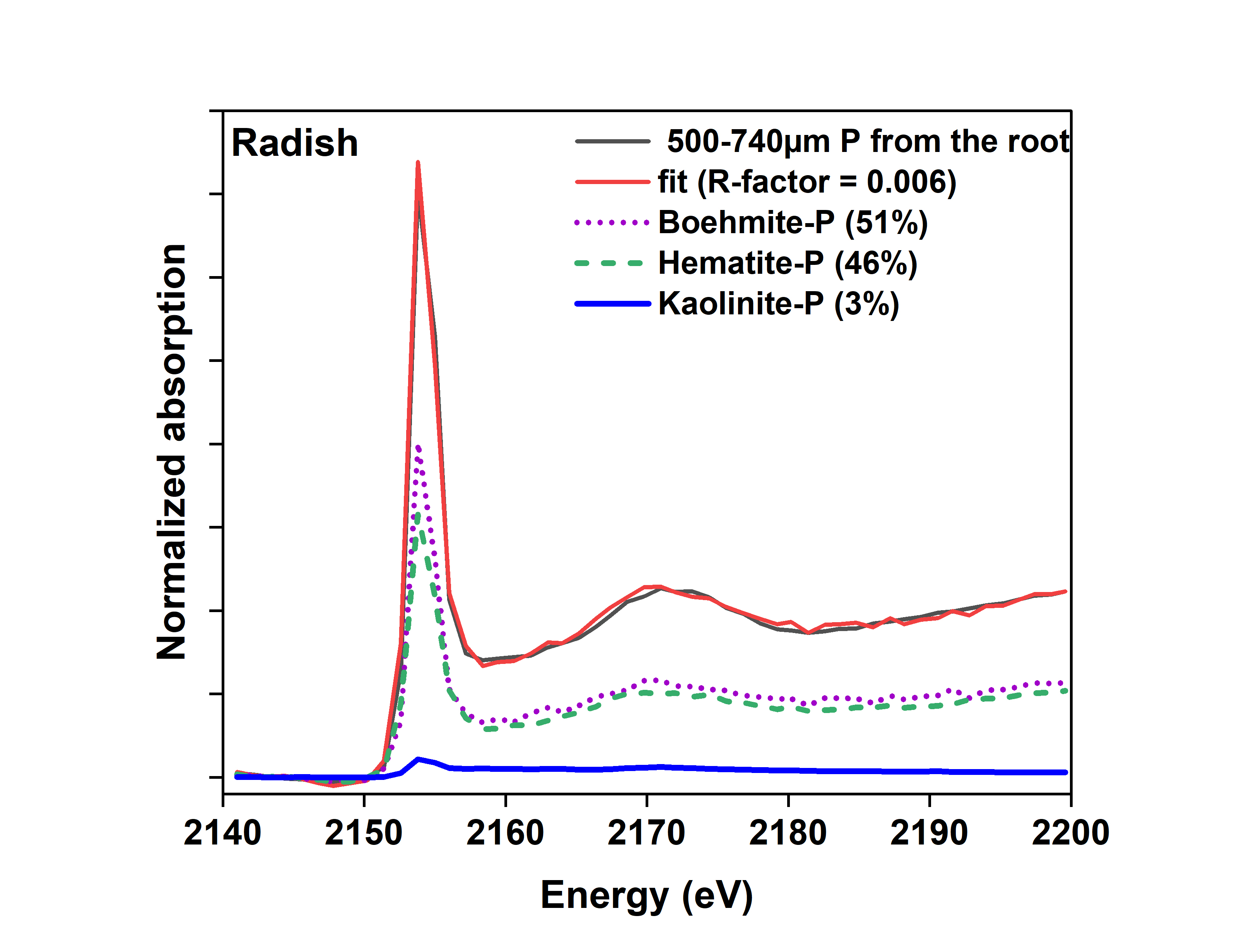

E

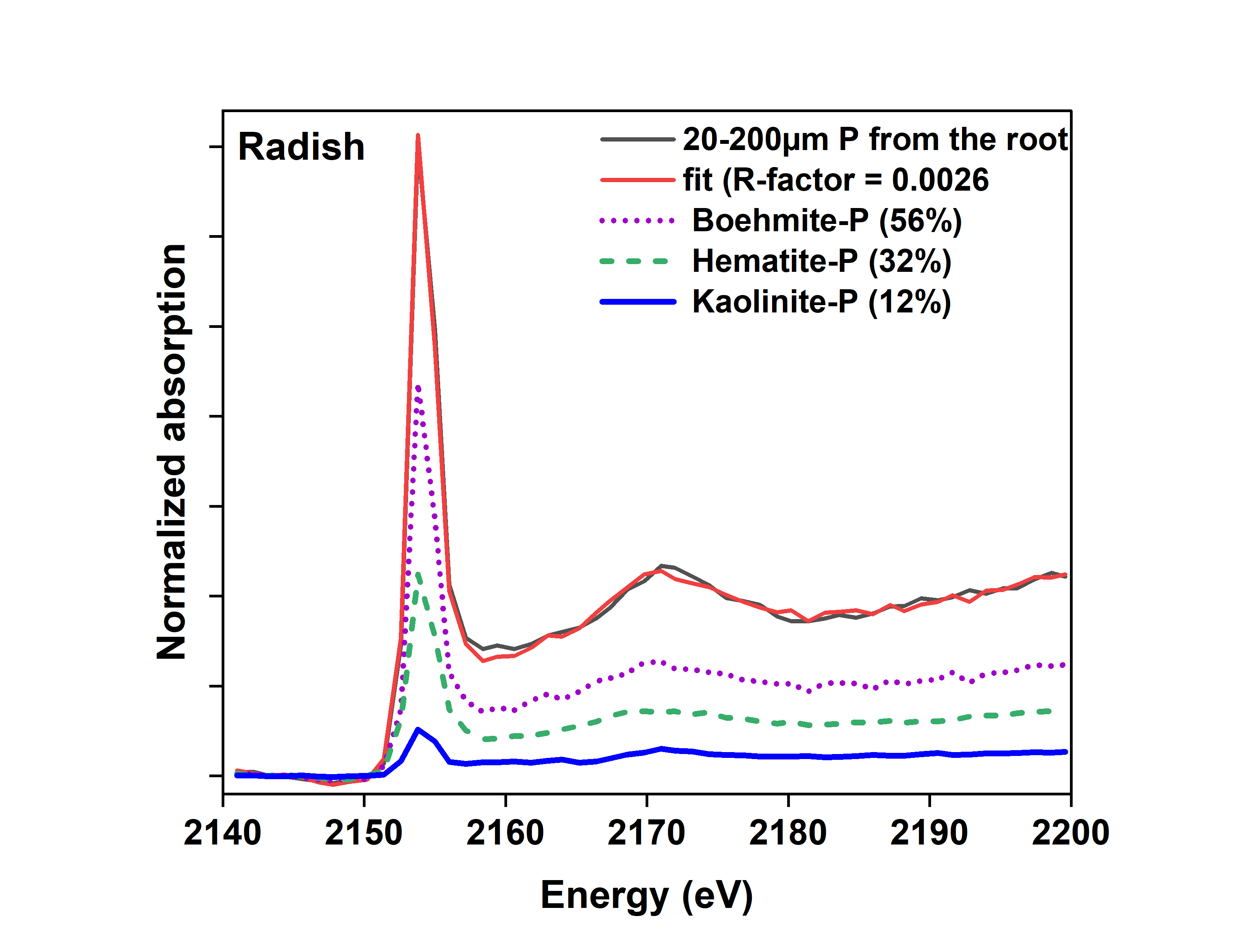

C

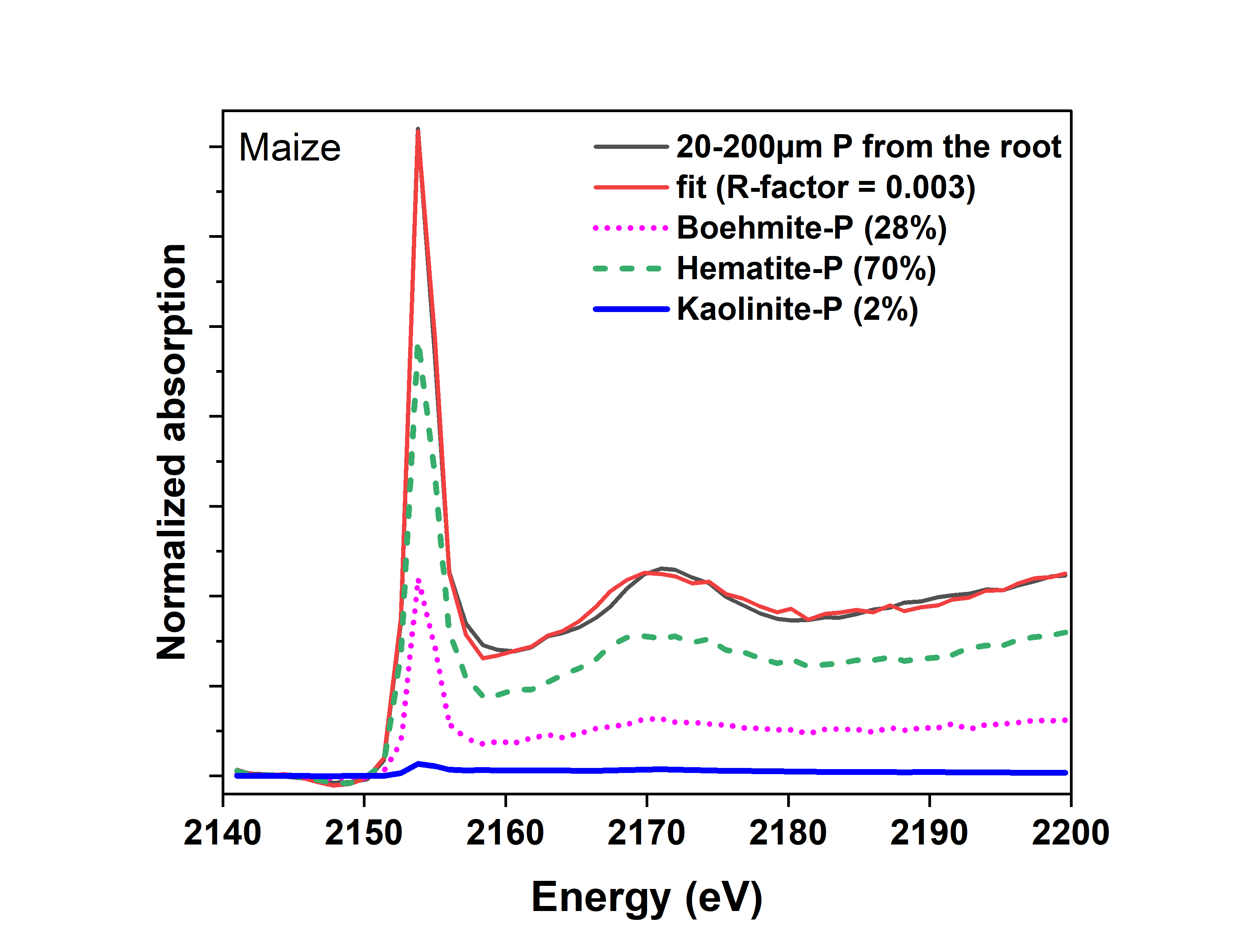

D

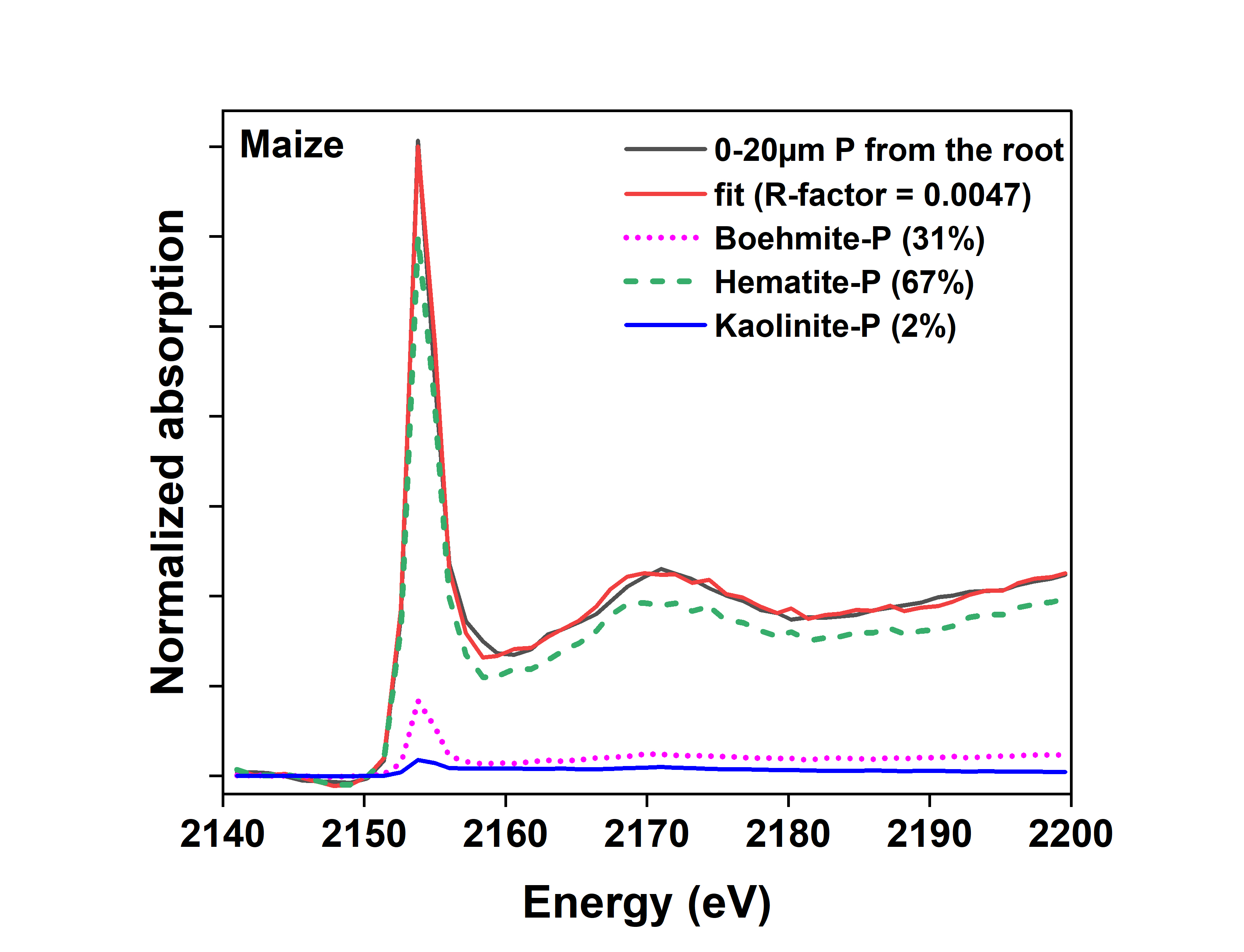

B

**Figure S7:** Normalized Phosphorus K-edge XANES spectra of each cluster group from the PCA (A) Radish 0-20µm from the root (B) Maize 0-20µm from the root, (C) Radish 20-200µm from the root, (D) Maize 20-200µm from the root, (E) Radish 500-740µm from the root (F) Maize 500-740µm from the root. Values in the parentheses indicate percentage of P species calculated by linear combination fitting

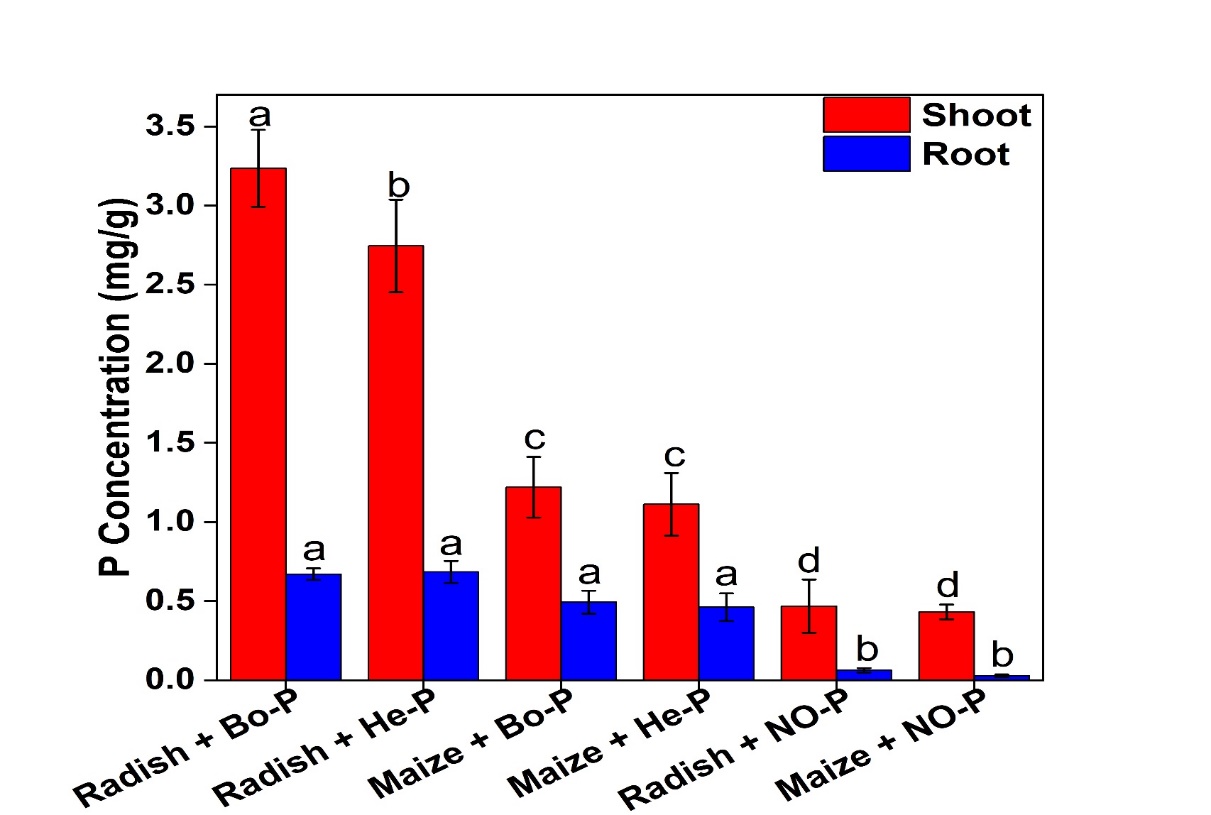

**Figure S8:** Total phosphorus concentration (mg/g) in shoot and root of radish and maize. Radish +Bo-P and Maize +Bo-P are radish and maize plants respectively, planted in substrate contains mixture of pure sand, strongly adsorbed P on boehmite (25mg P), kaolinite and irrigated with nutrient solution without P. Radish + He-P and Maize + He-P are radish and maize plants respectively, planted in substrate contains mixture of pure sand, strongly adsorbed P on hematite (25mg P), kaolinite and irrigated with nutrient solution without P. Radish + NO-P and Maize + NO-P (Control) are radish and maize plant respectively planted in a substrate contains mixture of pure sand, pure hematite, pure boehmite, and pure kaolinite irrigated with nutrient solution without P. The error bars indicated the standard deviation of replicates (n=5). The letters a, b, c, and d on the bars indicate significant differences between treatments. Treatments with the same letter shows that the treatments are not significantly different, while different letters signify significant differences (P < 0.05).
